# Tracing the *In Vivo* Fate of Nanoparticles with a “Non-Self” Biological Identity

**DOI:** 10.1101/2020.03.27.012146

**Authors:** Hossein Mohammad-Beigi, Carsten Scavenius, Pia Bomholt Jensen, Kasper Kjaer-Sorensen, Claus Oxvig, Thomas Boesen, Jan J. Enghild, Duncan S. Sutherland, Yuya Hayashi

## Abstract

Nanoparticles can acquire a biomolecular corona with a species-specific biological identity. However, “non-self” incompatibility of recipient biological systems is often not considered, for example, when rodents are used as a model organism for preclinical studies of biomolecule-inspired nanomedicines. Using zebrafish embryos as an emerging model for nano-bioimaging, here we unraveled the *in vivo* fate of intravenously injected 70 nm SiO_2_ nanoparticles with a protein corona pre-formed from fetal bovine serum (FBS), representing a non-self biological identity. Strikingly rapid sequestration and endolysosomal acidification of nanoparticles with the pre-formed FBS corona were observed in scavenger endothelial cells within minutes after injection. This led to loss of blood vessel integrity and inflammatory activation of macrophages over the course of several hours. As unmodified nanoparticles or the equivalent dose of FBS proteins alone failed to induce the observed pathophysiology, this signifies how the corona enriched with a differential repertoire of proteins can determine the fate of the nanoparticles *in vivo*. Our findings thus reveal the adverse outcome triggered by incompatible protein coronas and indicate a potential pitfall in the use of mismatched species combinations during nanomedicine development.

---

Biomimetic camouflage by exploitation of “self” molecules presented at nanoparticles is a strategy for prolonging the circulation half-life to target tumors *in vivo.^1–3^* Self biomolecules, when they adsorb to nanoparticles *in vivo* in a natural but non-specific manner^3–6^ in contrast to a pre-designed pattern, constitute a biological identity known as a biomolecular corona.^7^ If the presented repertoire consists partially/entirely of proteins from a different species, the protein corona (PC) is then considered “non-self” rather than camouflage. Unlike adaptive immunity that tolerates self molecules to detect any antigens other than self, non-self detection in innate immunity relies on the complement system and pattern recognition of conserved motifs associated with pathogens/danger signals at the cellular level.^8^ The non-self PC could thus be interpreted as local enrichment of potentially immunogenic (foreign) proteins^9^ presenting molecular patterns with or without conserved domains and conformations. Despite the prevailing idea of the biological identity, fetal bovine serum (FBS) is still routinely used as a cell culture supplement for testing the toxicity/biocompatibility of nanoparticles in non-bovine cell lines anticipating *in vivo* translation of the knowledge. Only a few *in vitro* studies exist, however, that concerned the mismatched combinations of the species origin of the cell lines and nanoparticles’ species identity with a proper control.^10–13^ To our knowledge, no one has addressed how the nonself biological identity determines the fate of the nanoparticles *in vivo*.

Zebrafish embryos have recently emerged as a bioimaging model for preclinical screening of nanomedicines^14–22^ with an intriguing opportunity of whole-embryo fixation for transmission electron microscopy (TEM) to visualize intracellular trafficking of nanoparticles *in vivo* at high resolution.^23, 24^ Here we used the intravital real-time/ultrastructural imaging approaches established previously^24^ and the basic technical and biological knowledge are described therein. Of note is the scavenger phenotype of venous endothelial cells (ECs) that lines the caudal vein (CV) and the CV plexus, the functional analogue of liver sinusoidal ECs in mammals.^15^ They sequester blood-borne SiO_2_ nanoparticles in a scavenger receptor-dependent manner^24^ with blood clearance kinetics resembling that of mammalian models.^25^ In this study, 70 nm SiO_2_ nanoparticles were intravenously (IV) injected with or without pre-formed FBS PC. The assumption here is that bare nanoparticles, like in other *in vivo* experimentation, acquire a self biological identity as soon as they are introduced to the bloodstream,^26^ while pre-formed FBS PC retains its compositional profile as a non-self biological identity until localized intracellularly where degradation may take place.^27, 28^ The former condition is hereafter referred to as “unmodified nanoparticles” in contrast to the latter condition “FBS-PC nanoparticles”. By tracing the *in vivo* fate of the non-self biological identity, we here find rapid nanoparticle sequestration by scavenger ECs within a time-scale of minutes post-injection (mpi) and strong induction of pro-inflammatory responses in macrophages that last several hours. This indicates a clear difference in the mechanism and kinetics of nanoparticle sequestration triggered by the preformed FBS PC with a previously undescribed potential of a non-self biological identity to elicit inflammation *in vivo*.

## RESULTS AND DISCUSSION

### Exposure to blood plasma proteins does not overwrite but adds to pre-formed coronas

A critical premise for testing non-self biological identities *in vivo* is the retention of the preformed FBS PC until eventually cleared from the bloodstream *via* ligand-receptor interactions at the cell surface. We started with characterizing 70 nm SiO_2_ nanoparticles of three different surface types (Plain or functionalized either with -NH_2_ or -COOH) with or without pre-formed FBS PC. The SiO_2_ nanoparticles represent a model system especially suited for this study owing to the matrix-incorporated fluorescent tracers of choice, intrinsic high electron density of SiO_2_, and the large particle size that allows excellent recovery of the nanoparticle-protein complexes even after extensive washing by centrifugation. To form FBS PC, the nanoparticles were mixed in 90% (v/v) heat-inactivated FBS and washed by multi-step centrifugation as previously described.^13^ The FBS-PC nanoparticles displayed a colloidally-stable population (Figure 1a,b and Supporting Figure 1) with an increased hydrodynamic diameter of ~30 nm for the plain surface and ~50 nm for NH_2_ and COOH surfaces (Supporting Table 1).

**Figure 1.**
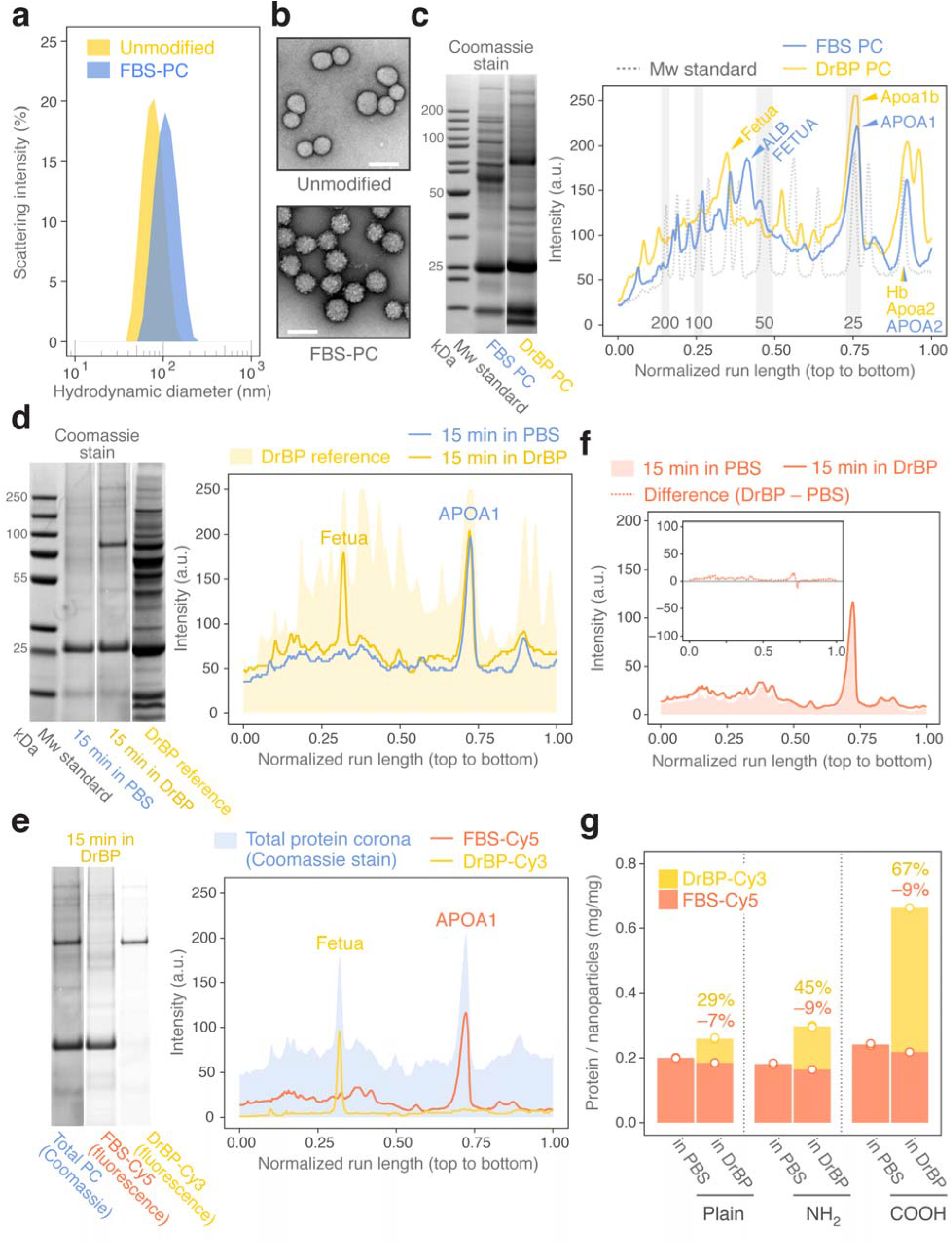
Characterization of FBS-PC nanoparticles and retention of the PC *in vitro*. (a) DLS analysis on the hydrodynamic size distribution verifying a stable dispersion of FBS-PC nanoparticles. (b) TEM images of negatively-stained 70 nm SiO_2_ nanoparticles. FBS-PC nanoparticles show the globular appearance of dehydrated proteins at nanoparticles. Scale bars, 100 nm. (c,d) SDS-PAGE profiles of Coomassie Brilliant Blue-stained corona proteins (nominal mass of nanoparticles loaded: 150 μg in c, 30 μg in d). Intensity profiles are shown alongside the gel images. (c) FBS PC is compared to DrBP PC to reveal proteins with an intrinsically high affinity for the SiO_2_ nanoparticles. (d) Incubation of FBS-PC nanoparticles in DrBP (see “DrBP reference” for the total protein repertoire) adds further proteins to the corona. Selected bands were excised for protein identification and the peaks annotated in the profile plot, except for Apoa1 previously identified.^13^ Shaded peaks indicate proteins common to FBS and DrBP. Original gel images and details of identified proteins are shown in Supporting Figure 2. (e) Fluorescence scans distinguish the origins of the two highly enriched corona proteins, which were further validated by MS/MS protein identification. (f) SDS-PAGE profiles of FBS-Cy5 reveal no particular removal of FBS PC even following 15 min incubation in DrBP. (g) Fluorimetry-based quantification of FBS-Cy5 and DrBP-Cy3 in the PC. Two other surface types of 70 nm SiO_2_ nanoparticles are included (-NH_2_ and -COOH; see Supporting Figures 1-4 for all characterization results). The values shown above the columns are the percentage decrease of FBS as compared to “in PBS” and the fraction of DrBP in the total PC. Columns represent the mean of two measurements (empty points). a.u., arbitrary units.

To study the retention of FBS PC *in vitro*, we used zebrafish blood plasma (DrBP) harvested from adult male fish^13^ representing the *in vivo* blood protein repertoire. It should be noted, however, that the liver of the zebrafish embryo is immature at 3 days post-fertilization (dpf) and the plasma proteome in adult fish^29^ does not entirely reflect the native protein repertoire at this embryonic stage.^30^ To identify proteins with a high affinity for the nanoparticles, PC made purely of FBS or DrBP was analyzed by SDS-PAGE and tandem mass spectrometry (LC-MS/MS) for protein identification (Supporting Table 2 lists the identified proteins). While Fetua was specifically enriched in DrBP PC and ALB in FBS PC, Apoa1b/APOA1 and Apoa2/APOA2 were two common proteins that constitute a major component in both PC types (Figure 1c and Supporting Figure 2) confirming previous results.^13^ Notably, nanoparticles with the COOH surface showed PC profiles skewed towards high molecular weight proteins for both of FBS PC and DrBP PC, while those at the NH_2_ surface were similar to at plain nanoparticles (Supporting Figure 3). The observed differences in the pattern of corona proteins between FBS PC and DrBP PC are attributed primarily to the differential protein repertoires due to species differences, let alone the protein source: serum *versus* plasma. The plasma protein repertoires of mammals and zebrafish are well-conserved; 92% of the identified zebrafish plasma proteome could be mapped to unique human orthologues.^31^ Yet, zebrafish blood lacks serum albumin and additionally contains numerous protein duplicates.^31^ In this study, FBS PC was indeed partially characterized by the enrichment of serum albumin (ALB) that does not exist in DrBP PC. One of the apolipoprotein A1 duplicates specific to zebrafish, Apoa1b, also appeared as the counterpart to bovine APOA1. At the compositional level, therefore, these dissimilarities could contribute to the non-self biological identity of FBS-PC nanoparticles. This could be further complicated by the extent of species cross-reactivity between ligands (epitopes presented at the PC) and receptors, if one considers the structural, rather than functional, properties of each corona protein. Quantification of such “foreignness” is, however, not within the scope of this study and thus remains to be answered.

Next, we incubated FBS-PC nanoparticles in DrBP for a minimum of 15 min to characterize PC retention *in vitro*, corresponding to previously estimated minimal elimination half-life of 14 min following IV injection of the SiO_2_ nanoparticles but without pre-formed FBS PC.^24^ Fluorescence labeling and MS/MS approaches confirmed retention of APOA1 in the FBS PC while Fetua was added from the 15 min incubation in DrBP (Figure 1d,e and Supporting Figures 2 and 4). This was particularly prominent for the plain surface, with only 7% reduction in proteins of FBS origin and 29% additional proteins acquired from DrBP (Figure 1f,g). The other two surface types had 9% reduction of FBS PC especially in the high molecular weight range but attracted even greater amounts of DrBP representing close to or more than the half of the “mixed” species identity of the newly formed PC (Figure 1g, and Supporting Figures 2 and 4). These *in vitro* results need to be interpreted cautiously, because, in addition to the anticipated difference of the plasma proteomes, the spontaneous formation of PC *in vivo* tends to display more diverse arrays of proteins than the counterpart *ex vivo* PC as demonstrated for mice^5^ and humans.^6^ The *in vivo* milieu of the bloodstream, rich in biomolecules and other biological entities, could thus competitively replace a certain fraction of the pre-formed PC (as a non-self biological identity) as fast as 5 min post-injection in mice, although unclear in the quantification.^32^ Here our purpose was to maximally retain the introduced non-self biological identity of the pre-formed PC, so we selected plain surface nanoparticles with minimal addition of blood proteins *in vitro* for further studies. Interestingly, prolonged exposure to DrBP (6 h) resulted in a decrease of the Fetua content but an increase in the Apoa1b:APOA1 ratio indicating an exchange of the same type of proteins that differ in the species origin (Supporting Figures 2 and 5).

### Nanoparticles with pre-formed FBS protein corona are rapidly sequestered

Nanoparticles with or without pre-formed FBS PC were IV-injected into the common cardinal vein (CCV) of 3 dpf zebrafish embryos for systemic circulation *via* the heart to the peripheral vasculature. At 1 hour post-injection (hpi), while the majority of unmodified nanoparticles were still retained in the bloodstream, nearly complete blood clearance was already observed for FBS-PC nanoparticles with a biodistribution pattern specific to veins (Figure 2a and Supporting Figure 6). Those veins include the CV/CV plexus and to a lesser extent the CCV, posterior caudal vein (PCV), the intersegmental veins (SeV), the primary head sinus (PHS; a major venous drainage of cerebral veins) and the primordial midbrain channel (PMBC; the vessel connecting cerebral veins) indicating a higher probability of interactions with scavenger ECs (see Supporting Figure 6 for each anatomical annotation) as observed for anionic liposomes.^15^ To resolve the rapid sequestration kinetics, we then performed time-lapse imaging up to 30 mpi at the ROI indicated in Figure 2a (Movies 1 and 2). The nanoparticle concentration in the bloodstream was clearly lower for FBS-PC nanoparticles, as determined by the mean fluorescence intensity in the caudal artery (CA), with a faster exponential decay function compared to unmodified nanoparticles (Figure 2b-d). Nanoparticle sequestration was also strikingly rapid for FBS-PC nanoparticles initiated with massive adherence to scavenger ECs rather than to macrophages as observed for unmodified nanoparticles (Figure 2b,c,e).

**Figure 2.**
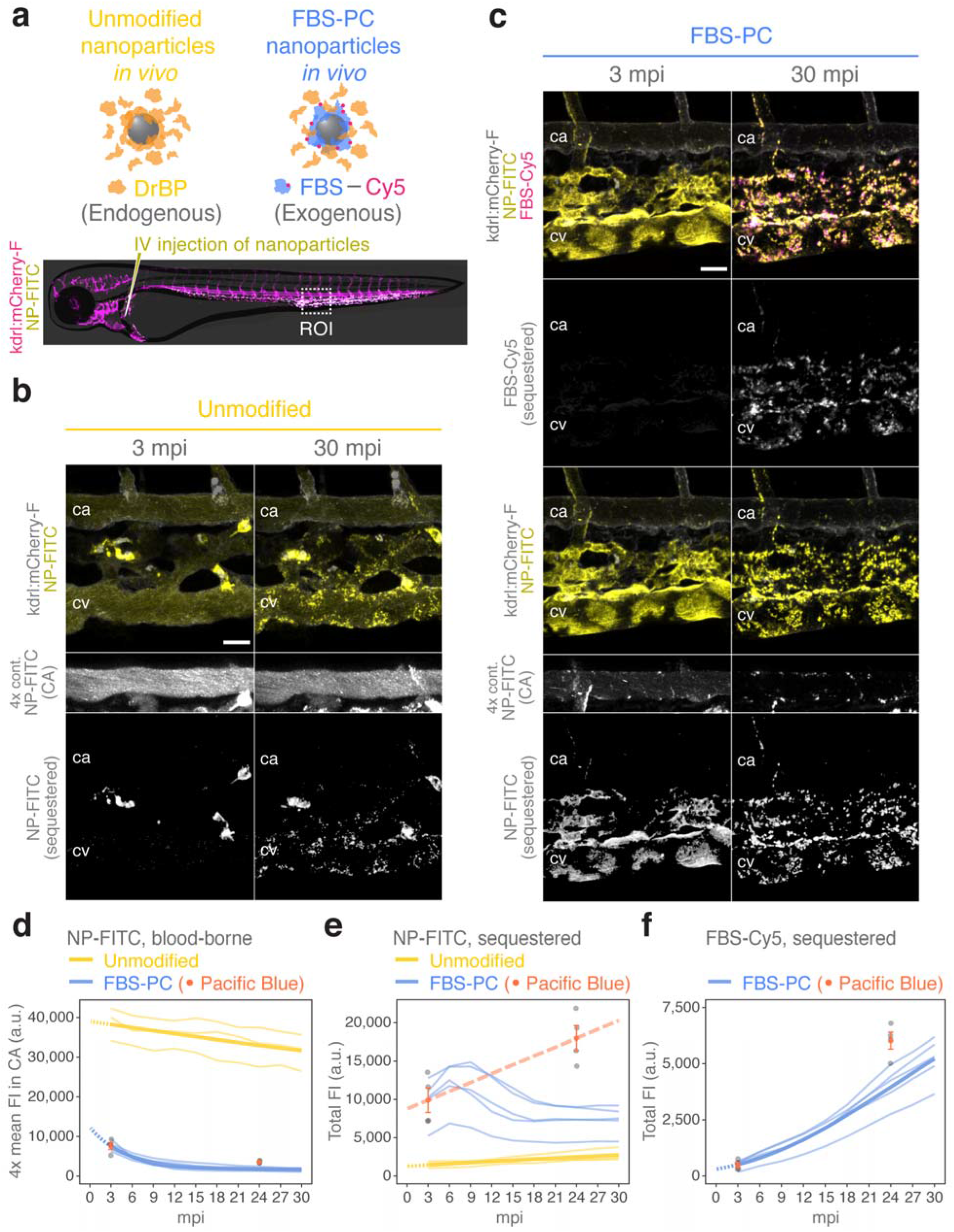
*In vivo* imaging of FBS-PC nanoparticle sequestration. (a) Schematics representing the unmodified and FBS-PC nanoparticles *in vivo*. The panel below depicts the blood vessel network in a 3 dpf embryo with an overlaid illustration of the whole body. See Supporting Figure 6 for the original images showing biodistribution of the injected nanoparticles. (b-f) *Tg(kdrl:mCherry-F)* embryos at 3 dpf were injected with FITC-labeled SiO_2_ nanoparticles (NP-FITC) with or without pre-formed FBS PC (FBS-Cy5) and imaged every 3 min. Representative images showing sequestration of (b) unmodified and (c) FBS-PC nanoparticles. The huge clusters of nanoparticles found in (b) are likely those associated with macrophages.^24^ The NP-FITC signals in the CA are multiplied by 4-fold to aid visualization. Anterior left, dorsal top. Scale bars, 20 μm. Kinetics for (d) clearance of blood-borne nanoparticles and sequestration of (e) nanoparticles and (f) FBS PC are plotted as individual embryos (thin lines, *n* = 3 for unmodified, *n* = 5 for FBS-PC) and, where possible, curve-fitted (thick lines). Dashed lines between 0-3 mpi are predictions from the fitted models. Results from Pacific Blue-labeled nanoparticles are plotted for the two-time points as individual embryos (gray points, *n* = 4) and as the mean ± SE (orange-colored). Note that the values are estimated from the relative difference between 3 and 24 mpi, where the mean values at 3 mpi are the same for FITC-labeled and Pacific Blue-labeled nanoparticles. Linear fitting (the long-dashed line) is shown in (e). a.u., arbitrary units. FI, fluorescence intensity. See Movies 1 and 2 for the time-lapse sequences.

The measured fluorescence intensity from sequestered FBS-PC nanoparticles (NP-FITC) in scavenger ECs, however, did not continue to increase (Figure 2c,e) despite the consistent accumulation of FBS-Cy5 representing FBS PC (Figure 2c,f). We attribute this lack of increase to pH-dependent quenching of the FITC and found that >50% of NP-FITC fluorescence was indeed lost at a pH mimicking the lysosome (~4.7)^33^ while FBS-Cy5 was little affected both as a label for free proteins and PC (Supporting Figure 7). Pacific Blue, on the other hand, has *pK_a_* of 3.7 and documented stability in acidic organelles in live cells.^34^ As this dye is prone to photobleaching, we opted for single-timepoint experiments on two sets of embryos at 3 and 24 mpi to calculate relative changes. FBS-PC nanoparticles (Pacific Blue) were thus prepared likewise to complement the FITC dataset. Estimated values based on the relative proportions are plotted along with the time-lapse results, validating the kinetics for blood clearance (Figure 2d) and FBS PC accumulation (Figure 2f) and indicating how the total fluorescence intensity for sequestered nanoparticles would have increased if FITC was not quenched (Figure 2e). FBS-PC nanoparticles were thus rapidly sequestered in scavenger ECs experiencing a local pH decrease at a faster rate than that of unmodified nanoparticles. The unmodified and FBS-PC nanoparticles also revealed a contrasting pattern in the cell types (macrophages *versus* scavenger ECs) that play a dominant role for membrane associations within minutes after IV injection.

### The protein corona undergoes acidification after endolysosomal sequestration

To verify that the loss of NP-FITC fluorescence was caused by endolysosomal acidification following nanoparticle uptake, we have devised a dual labeling strategy by which the fluorescence ratio of pHrodo (pH indicator) to Cy5 (pH-insensitive dye) responds to local acidification within a cell. The fluorescence response of the dyes as a label for free proteins and nanoparticle-protein complexes in the pH range relevant for endosomes (pH 5.5-6.3) and lysosomes (pH 4.7)^33^ confirmed the positive correlation between acidification and increase in the pHrodo/Cy5 ratio (Figure 3a and Supporting Figure 7). We then quantified the fluorescence ratio *in vivo* in zebrafish embryos using a 3D mask approach^24^ whereby the colocalization of Cy5 and pHrodo signals was determined at each focal plane of 2 μm-thick optical sections before maximum intensity projection of a stacked image. The increase in the pHrodo/Cy5 ratio was clearly visible between 6 to 12 mpi (Figure 3b and Movie 3) but the largest change was in fact estimated to be occurring already at 5 mpi (Figure 3c). This time-frame of ~10 min corresponds to the time it takes for newly formed endolysosomes to establish the lysosomal pH for acid hydrolase activity.^35^

**Figure 3.**
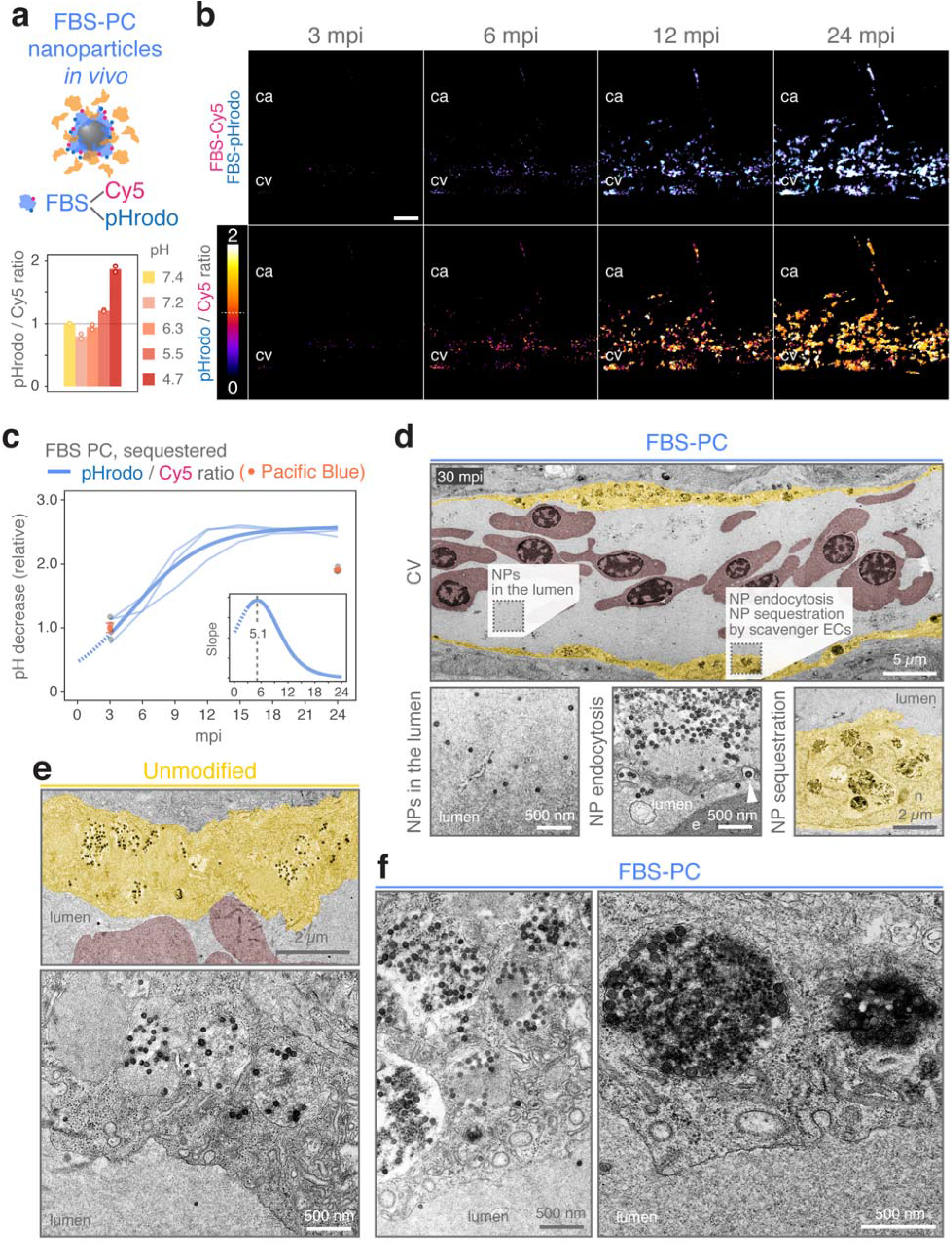
Acidification of FBS-PC nanoparticles following endolysosomal sequestration. (a) Schematic illustrating dual-labeled FBS-PC nanoparticles for quantification of pHrodo-to-Cy5 ratio. The graph below shows the fluorescence ratio *in vitro* measured by fluorimetry at the intracellular pH range (mean, *n* = 2), indicating an increase in the ratio at pH 5.5 and 4.7. (b,c) Wild-type embryos at 3 dpf were injected with FITC-labeled SiO_2_ nanoparticles with pre-formed FBS PC (FBS-Cy5/-pHrodo) and imaged every 3 min. (b) Representative images showing the fluorescence ratios, 0-2 scaled. Anterior left, dorsal top. Scale bars, 20 μm. (c) The median fluorescence ratios are plotted relative to 3 mpi as individual embryos (thin lines, *n* = 3) and curve-fitted (the thick line). The dashed line between 0-3 mpi is a prediction from the fitted model. The slope of the fitted curve is shown in the inset with the vertical dashed line indicating the peak maximum. Results from Pacific Blue-labeled nanoparticles are plotted for the two-time points as individual embryos (gray points, *n* = 4) and as the mean ± SE. (orange-colored). See Movie 3 for the time-lapse sequence. (d-f) Wild-type embryos at 3 dpf were injected with FITC-labeled SiO_2_ nanoparticles with or without pre-formed FBS PC and chemically-fixed at 30 mpi for TEM imaging. Representative (tiled) images indicating (d) the observed aspects of particular note, (e) sequestered unmodified nanoparticles and (f) FBS-PC nanoparticles found in highly electron-dense vesicular lumen. In low magnification images, scavenger ECs and erythrocytes are pseudo-colored in yellow and red, respectively, to visually aid the boundaries between the cells and the blood lumen. NP, nanoparticle. e, erythrocyte. n, nucleus.

At the ultrastructural level, FBS-PC nanoparticles were singly dispersed in the blood lumen, internalized *via* endocytosis and sequestered in endolysosomal compartments in scavenger ECs (Figure 3d) as were observed for unmodified nanoparticles (Figure 3e). The number of compartmentalized nanoparticles was clearly higher for FBS-PC nanoparticles (Figure 3f) in good agreement with the intravital imaging approach (Figure 2). The lumen of those endolysosomal compartments observed with FBS-PC nanoparticles differed in the electron density displaying light-to-dark contrast (Figure 3f), possibly due to acidification of the vesicle.^36^ FBS-PC nanoparticles thus undergo rapid cellular sequestration and subsequently experience acidification in the endolysosomes in scavenger ECs. These biological processes we observed here mostly occurred within the first 30 mpi, which underscores the importance of resolving early phase kinetics in the *in vivo* setting where rapid blood clearance is frequently reported also in mammalian models. ^3, 21, 25, 37^ Within the specimens at this chosen time point (30 mpi), however, we were unable to identify macrophages that display signatures of nanoparticle uptake. This is probably because the sequestration of FBS-PC nanoparticles in macrophages was much less pronounced than that of unmodified nanoparticles as indicated by intravital imaging at 3-30 mpi (Figure 2b,c).

### The protein corona is degraded over time concurrently with loss of blood vessel integrity and induction of pro-inflammatory responses

As FBS-PC nanoparticles were confined in the endolysosomal pathways, we next determined degradation of the PC by quantifying the fluorescence ratio of FBS-Cy5 to NP-Pacific Blue at selected time-points (1, 2, 4 and 6 hpi). To assess the ratio separately for ECs and macrophages, double transgenic *Tg(fli1a:EGFP); Tg(mpeg1:mCherry)* embryos were used in conjunction with the 3D mask approach for colocalization of FBS PC nanoparticle signals with cell types of interest (Supporting Figure 8). The FBS-Cy5 to NP-Pacific Blue ratio was significantly lower for macrophages at 6 hpi (60%) and to a lesser extent for ECs (79%), suggesting PC degradation to become prominent at between 4 and 6 hpi (Figure 4a-d). All other parameters tested were not significantly different except for an unexpected lower cell area of ECs at 1 hpi (Supporting Figure 9). With a similar time-course, earlier *in vitro* studies reported lysosomal degradation of PC to occur around 7 h of exposure.^27, 28^

**Figure 4.**
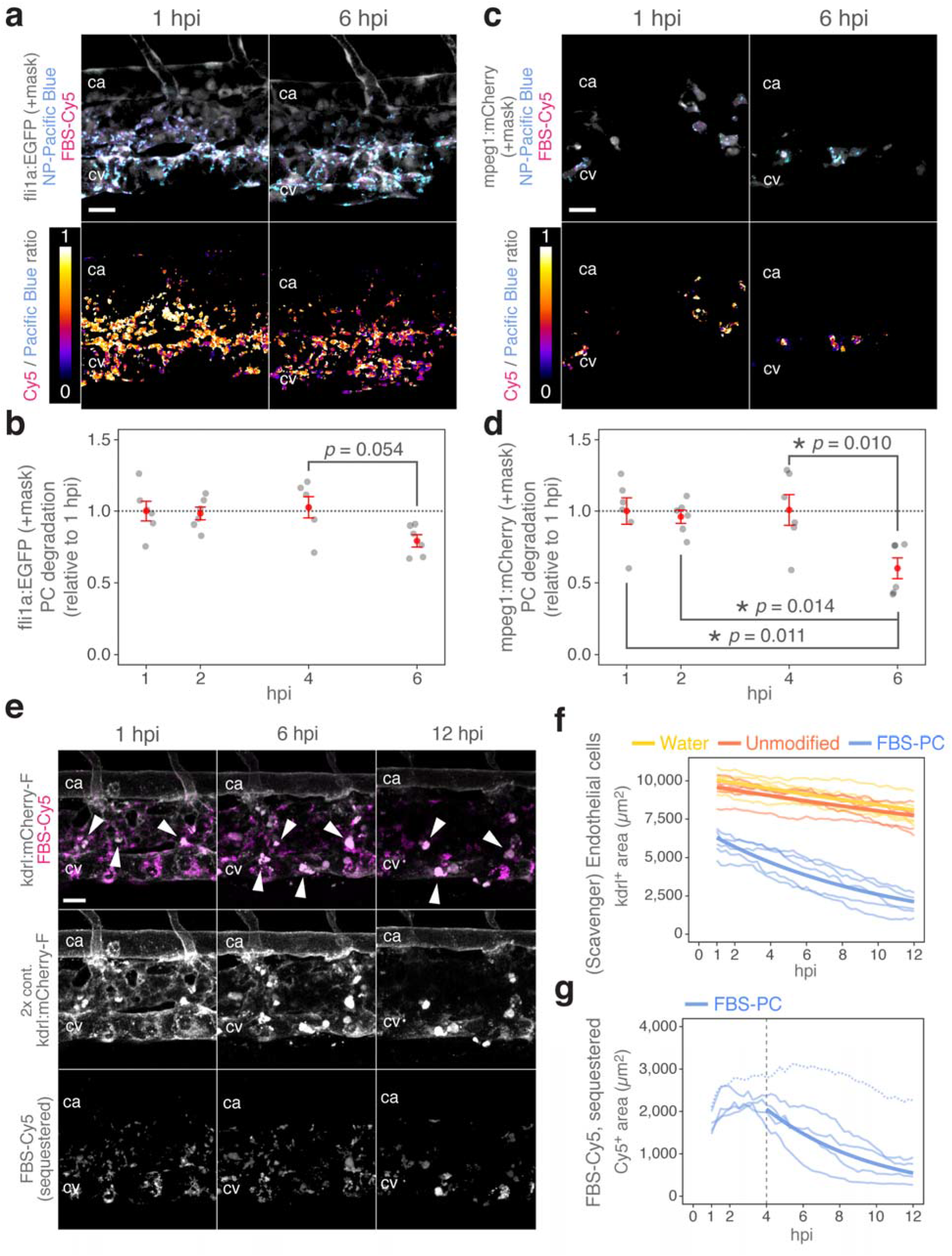
Degradation of PC and loss of blood vessel integrity. (a-d) *Tg(fli1a:EGFP); Tg(mpeg1:mCherry)* embryos at 3 dpf were injected with Pacific Blue-labeled SiO_2_ nanoparticles (NP-Pacific Blue) with pre-formed FBS PC (FBS-Cy5) and imaged independently at each time point. Representative images showing (a) EC- and (c) macrophage-specific signals for the Cy5-to-Pacific Blue ratios, 0-1 scaled. The median fluorescence ratios were plotted relative to 1 hpi for (b) ECs and (d) macrophages as individual embryos (gray points, *n* = 6) and the mean ± SE (red-colored). Significant differences were tested by one-way ANOVA with Tukey’s HSD post-hoc comparisons (degrees of freedom = 20, *F* values = 3.295 for ECs and 6.066 for macrophages). (e-g) *Tg(kdrl:mCherry-F)* embryos at 3 dpf were injected with FITC-labeled SiO_2_ nanoparticles with or without pre-formed FBS PC (FBS-Cy5) and imaged every 15 min starting at 1 hpi. Representative images showing loss of fluorescence signals from scavenger ECs and sequestered FBS PC. The mCherry signals are multiplied by 2-fold to aid visualization. Arrowheads denote blob-like mCherry signals (and to some extent, of FBS-Cy5) likely specific to sequestration in macrophages. Areas representing (f) ECs (the mCherry signals) and (g) sequestered FBS PC (the Cy5 signals) are plotted over time as individual embryos (thin lines, *n* = 5) and curve-fitted (thick lines). The curve fitting in (g) was performed after excluding an outlier (the dashed line) and between 4-12 hpi where steady decreases are observed. Anterior left, dorsal top. Scale bars, 20 μm. See Movies 4-6 for the time-lapse sequences.

This time-frame of 4-6 hpi corresponds to the initiation of FBS-Cy5 fluorescence decay observed in 12 h time-lapse imaging, delivering an estimate of overall 27% reduction from 4 to 6 hpi (Supporting Figure 10). However, we have noticed that the constitutive reporter signals labeling mainly ECs (*kdrl:mCherry-F*) consistently decreased over time towards 12 hpi of FBS-PC nanoparticles which was much less obvious for embryos injected with water (a vehicle control) or unmodified nanoparticles with a 5-times lower decay constant than that for FBS-PC nanoparticles (Figure 4e,f, Movies 4-6 and Supporting Figure 10). In particular, scavenger phenotype of ECs forming the blood vessels in the CV and CV plexus was clearly affected in contrast to ECs lining the CA. This shows a striking similarity to the gradual loss of *kdrl:GFP* signals in scavenger ECs passively targeted by liposomal delivery of a cytotoxic drug resulting in compromised blood vessel integrity.^15^ Underscoring the targeted cytotoxicity, the signals from sequestered FBS-PC nanoparticles also diminished along with the fading *kdrl:mCherry-F* signals (Figure 4g and Supporting Figure 10), likely an indication for the disintegrated vasculature.

In the subsequent experiment, we used double transgenic *Tg(mpeg1:mCherry); Tg(tnfa:EGFP-F)* embryos to determine the pathophysiological consequence by the inducible reporter for transcriptional activation of the pro-inflammatory cytokine *tnfa (tumor necrosis factor-alpha).^38^* We first noted an overall decrease in the total cell area of macrophages which was not treatment-specific, suggesting no particular recruitment of macrophages from other parts of the body (Figure 5a,b). We then quantified the macrophage-specific contribution to the overall FBS PC signals, as the scavenger ECs were critically affected losing their function as the predominant sink of sequestered FBS-PC nanoparticles (Figure 4e-g). The increasing contribution of macrophages to the overall FBS PC signals (Figure 5a,c) may explain, although speculative as yet, sweeping of sequestered FBS-PC nanoparticles that once belonged to scavenger ECs through phagocytosis of apoptotic/necrotic cell debris.^39^ As reported elsewhere,^40^ the blob-like *kdrl:mCherry-F* signals seen in Figure 4e represent EC-derived extracellular vesicles (e.g. apoptotic bodies) accumulated by macrophages, and some of the FBS-Cy5 signals indeed began to associate with them displaying a wandering behavior (note the highly mobile, blob-like *kdrl:mCherry-F* signals in Movie 6).

**Figure 5.**
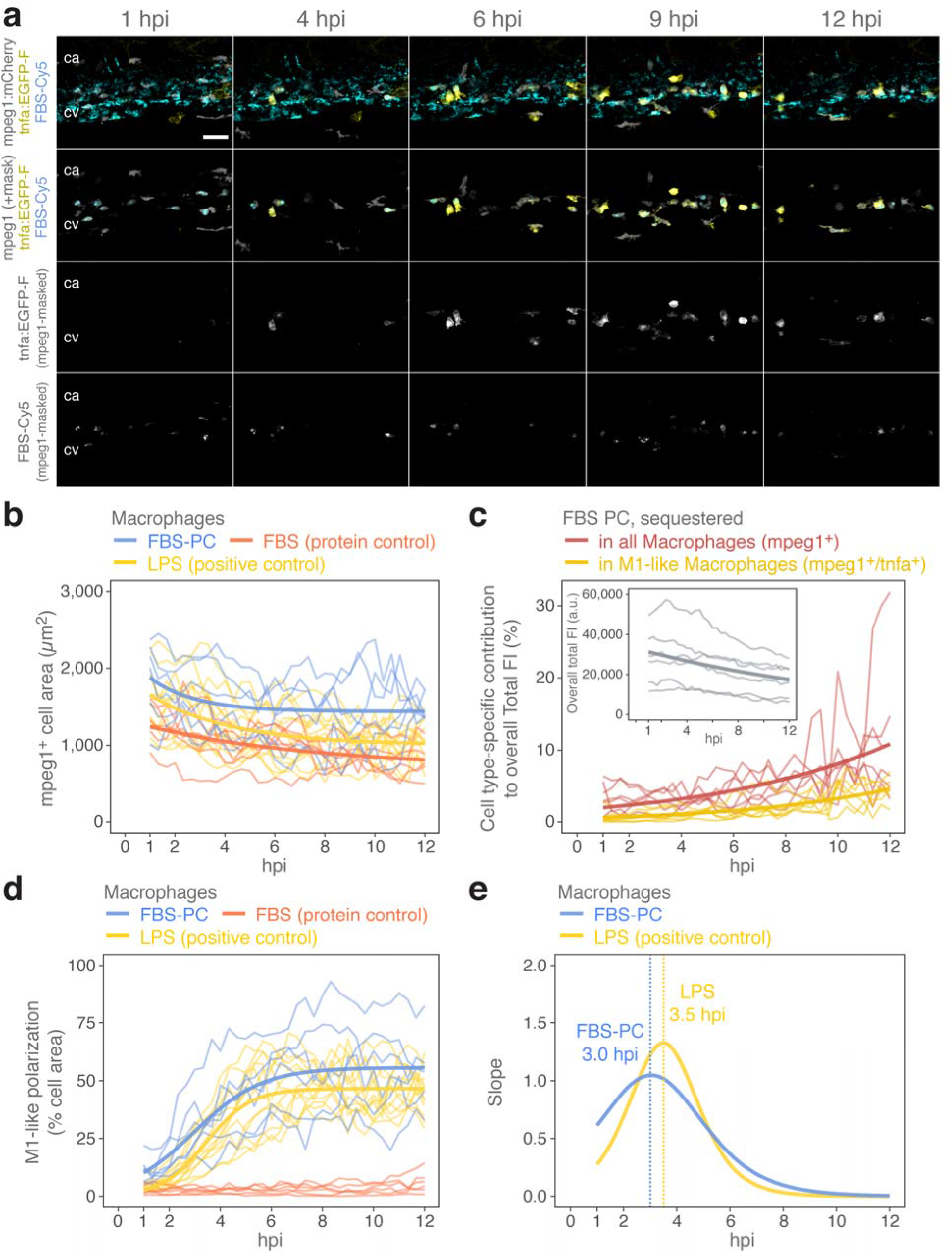
FBS-PC nanoparticles induce M1-like polarization of macrophages. (a-e) *Tg(mpeg1:mCherry); Tg(tnfa:EGFP-F)* embryos at 3 dpf were injected with Pacific Blue-labeled SiO_2_ nanoparticles with pre-formed FBS PC (FBS-Cy5), free FBS proteins only (at the equivalent protein mass of FBS-PC nanoparticles, 2 ng) or LPS (2.4 ng) and imaged every 20 min starting from 1 hpi. Representative images showing the time-course of FBS PC sequestration and induction of *tnfa* with or without the macrophage mask. Anterior left, dorsal top. Scale bars, 30 μm. Image analysis results and fitted models are plotted as individual embryos (thin lines, *n* = 6 for FBS-PC nanoparticles and FBS proteins only, *n* = 12 for LPS) and, where possible, curve-fitted (thick lines). Kinetics are shown for (b) the cell area of macrophages, (c) cell type-specific contributions to overall sequestration of FBS PC, (d) the degree of M1-like polarization represented by the percentage of colocalized areas between *tnfa^+^* and *mpeg1^+^*, and (e) the slopes derived from (d) with vertical dashed lines indicating the peak maxima. a.u., arbitrary units. FI, fluorescence intensity. See Movies 7-11 for the time-lapse sequences including negative results obtained from water control and unmodified nanoparticles.

Induction of *tnfa* is a molecular signature of M1-like (inflammatory) polarization in macrophages,^38^ and this phenotype propagated to ~50% (cell area) of macrophages within the ROI until approximately 6 hpi and persisted at 12 hpi (Figure 5d). This was observed only for FBS-PC nanoparticles and the positive control (LPS; lipopolysaccharides), and not for free FBS proteins at the equivalent dose, unmodified nanoparticles or water (Movies 7-11). The rate of propagation peaked at 3 hpi for FBS-PC nanoparticles and 3.5 hpi for LPS (Figure 5e), implying the involvement of different stimuli for the activation. It remains unclear, however, whether the pro-inflammatory cascade was initiated by macrophages or scavenger ECs. LPS-induced inflammation in zebrafish requires myeloid differentiation factor 88 (an adaptor for Toll-like receptor signaling axis)^41^ but the precise mechanism is yet uncertain unlike in mammals. In the case of FBS-PC nanoparticles, direct recognition of the non-self biological identity presented at the nanoparticles is an intriguing possibility, which could in principle be supported by a positive correlation between the FBS PC sequestration in macrophages and the pro-inflammatory responses. However, it seems unlikely the case as the M1-like phenotype-specific contribution to the sequestered FBS-Cy5 signals did not take over the contribution from non-polarized macrophages (Figure 5c) and those *tnfa^+^* macrophages without FBS-PC nanoparticles were also identified (Figure 5a and Movie 10). In contrast, our observation of scavenger EC disintegration at a corresponding time-course (Figure 4) favors a scenario in which extracellular pro-inflammatory stimuli such as damage-associated molecular patterns are involved in this pathophysiological consequence.^42, 43^

## CONCLUSION

Here we demonstrated the biological fate of FBS-PC nanoparticles and pathophysiological consequences *in vivo* using multicolor combinations of fluorescent tracers and endogenous reporters labeling ECs, macrophages and their M1-like phenotype in zebrafish embryos. To gain a holistic insight into the *in vivo* journey of the FBS PC, we addressed four aspects covering retention, sequestration, acidification and intracellular degradation of the PC that eventually lead to pro-inflammatory responses (Figure 6). The sequestration kinetics of the FBS-PC nanoparticles was strikingly different from that of unmodified nanoparticles whereby scavenger ECs likely play the predominant role in rapid clearance from the bloodstream. The scavenger ECs were thus heavily loaded with FBS-PC nanoparticles and as a consequence the blood vasculature was disintegrated, allowing macrophages to scavenge the cell debris along with sequestered nanoparticles. The inflammatory conditions are further supported by concurrent M1-like polarization of those macrophages with a time-frame of several hours after IV injection. This cannot be explained solely by the immune intolerance to the non-native protein repertoire presented by FBS, as the free proteins at the equivalent dose failed to mimic the pro-inflammatory responses. Our findings thus provide experimental *in vivo* evidence of the founding concept that the cell “sees” the biological identity^44^ to determine a differential fate of the same nanoparticles, visualized both in real-time and at ultrastructural resolution. This underscores the critical importance to carefully consider the species compatibility/extrapolation issues during preclinical *in vitro* and *in vivo* testing for the development of biomolecule-inspired nanomaterials.

**Figure 6.**
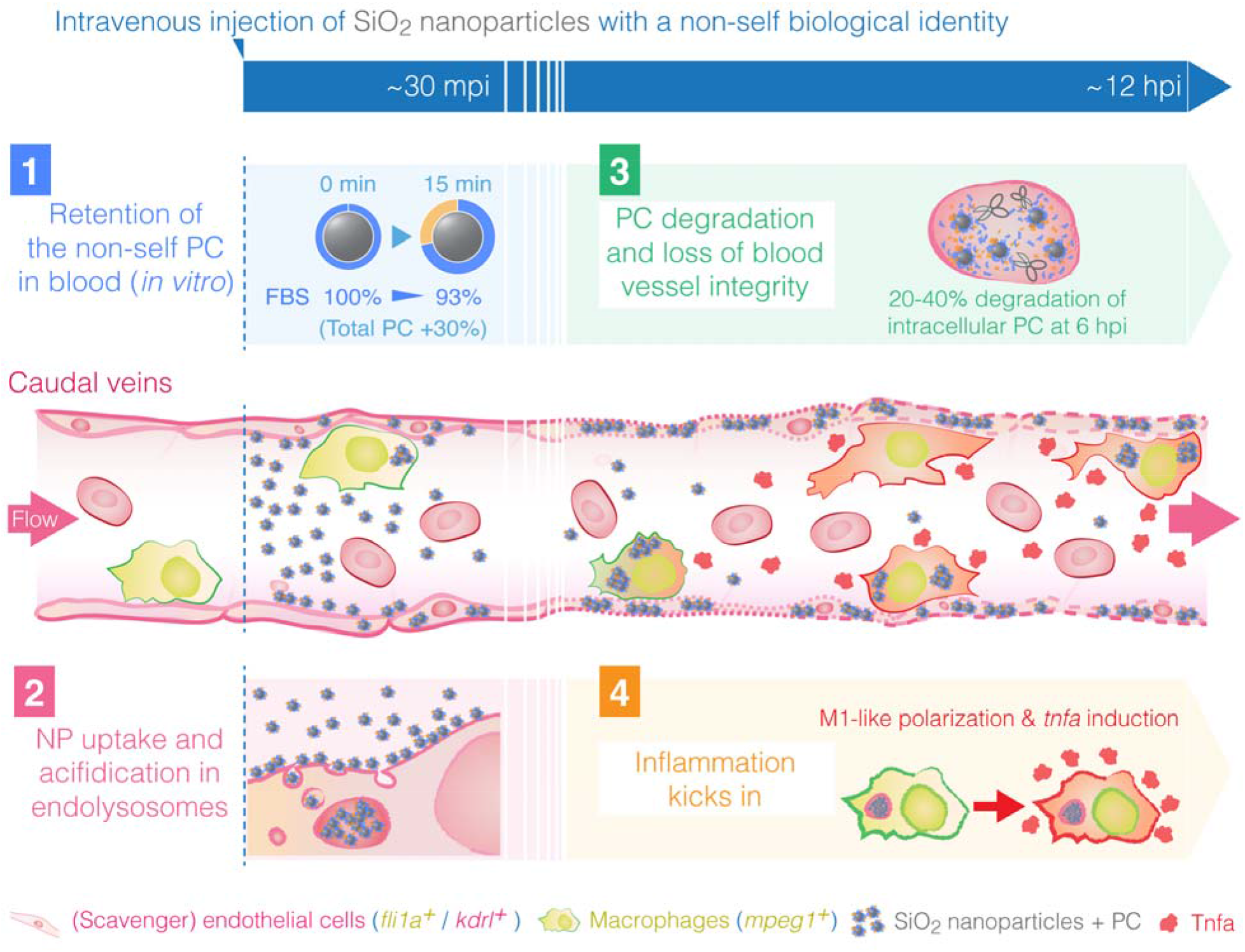
Schematic illustrating four elements directing the biological fate of nanoparticles with a non-self biological identity. (1) SiO_2_ nanoparticles may retain the pre-formed FBS PC as a non-self biological identity even after exposure to zebrafish blood plasma, however, with additional proteins that have a high affinity for the nanoparticles. (2) Within 30 min following IV injections, FBS-PC nanoparticles are rapidly sequestered by scavenger ECs and acidified in the endolysosomal compartments. (3) In a longer time-frame, degradation of the FBS PC occurs around 4-6 hpi in both scavenger ECs and macrophages while the former loses its integrity. (4) Concurrently, macrophages are activated to an inflammatory phenotype (M1-like polarization) that secretes the cytokine Tnfa coordinating the onset of inflammation.

## METHODS

### Preparation and characterization of FBS-PC nanoparticles

Fluorescently-labeled 70 nm SiO_2_ nanoparticles (plain, carboxyl and amine surfaces) were purchased from micromod Partikeltechnologie GmbH (Germany) and characterized by dynamic light scattering (DLS) and zeta potential measurements as described previously.^13^ Heat-inactivated FBS (ThermoFisher Scientific) was centrifuged at 16,000g for 3 min to remove any insoluble aggregates and the supernatant was mixed with nanoparticles at final concentrations of 90% (v/v) FBS and 0.4 mg/ml nanoparticles. Following 2 h incubation in darkness at 37°C, the nanoparticle-protein complexes were pelleted by centrifugation (20,000g, 20 min), PBS-washed three times, and redispersed in PBS at a nominal concentration of 4 mg/ml. The actual concentration was adjusted by fluorimetry using unmodified nanoparticles as the reference on a LS55 luminescence spectrometer (Perkin Elmer), where necessary, after direct labelling of PC as described below. The endotoxin level of FBS-PC nanoparticles was tested using a Pierce LAL Chromogenic Endotoxin Quantitation kit (ThermoFisher Scientific) and was 1.85 ± 0.07 EU/ml (mean ± SD of two independently prepared batches). The mass of proteins on nanoparticles was directly quantified using a Pierce BCA Protein Assay kit (ThermoFisher Scientific) with an additional centrifugation step (20,000g, 30 min) to collect the nanoparticle-free supernatant for absorbance measurements using a Varioscan plate reader (ThermoFisher Scientific). For TEM, specimens for unmodified and FBS-PC nanoparticles were prepared as previously described^13^ except that for negative staining uranyl formate was used instead of uranyl acetate.

### Fluorescence labeling of proteins

Proteins were in all cases labeled *via* the amine-reactive crosslinker N-hydroxysuccinimide (NHS) esters, available as water-soluble Sulpho-Cyanine 5 or Sulpho-Cyanine 3 NHS esters (Lumiprobe), or pHrodo Red NHS esters (ThermoFisher Scientific). For direct conjugation to corona proteins, 0.05 mg/ml of each dye (singly or as a mixture of Cyanine 5 and pHrodo Red) were incubated with FBS-PC nanoparticles for 2 h at room temperature (RT, 21°C). The reaction was quenched by 50 mM Tris, followed by PBS washing (20,000g, 20 min, three times centrifugation) to remove the free dyes. Free proteins were labeled following the manufacturer’s instructions and purified using Sephadex G-25 in PD-10 Desalting Columns (GE Healthcare Life Sciences).

### Fluorescence measurements at intracellular pH

The fluorescence intensity of unmodified nanoparticles, free proteins-labeled with dyes, and their complexes (FBS-PC nanoparticles) at pH relevant for intracellular environments was measured in phosphate-citrate buffer by fluorimetry as above. The nanoparticles did not show any signs of agglomeration between the pH range from 4.0 to 7.4, as determined by DLS.

### PC retention experiments *in vitro*

Collection of zebrafish blood plasma (DrBP) is described in full details in our earlier report.^13^. As DrBP obtained from adult female fish contains the yolk precursor proteins vitellogenins,^13^ for this study only male fish were chosen for the blood collection. To study the hard corona compositions, nanoparticles with FBS PC or DrBP PC (0.4 mg/ml) were washed by centrifugation (three times in PBS, 20,000*g*, 20 min) for SDS-PAGE. For the PC retention experiments, DrBP was Cy3-labeled as free proteins. Cy5-labeled FBS-PC nanoparticles (0.2 mg/ml) were incubated with 1 mg/ml Cy3-labeled DrBP (or only in PBS for a control) at 28°C (physiological temperature for zebrafish) in the dark for 15 min or 6 h, before proceeding to the centrifugation-based isolation for fluorimetry and SDS-PAGE. Protein mass was calculated based on known protein concentrations (BCA assays directly on FBS PC or on free FBS proteins) and fluorescence measurements by fluorimetry using reference unmodified nanoparticles (to estimate nanoparticle concentrations) and free FBS-Cy3 proteins (to estimate the mass of DrBP proteins added to PC.

### SDS-PAGE and gel scanning

Proteins bound to the nanoparticles were stripped by boiling for 5 min after the addition of 5× concentrated reducing sample buffer (5 % SDS, 50% glycerol, 100 mM DTT in 0.3 M Tris). The nanoparticle-free supernatant after centrifugation (20,000g, 30 min) was loaded on a Bolt 4-12% Bis-Tris Plus Gel at the volume corresponding to 150 μg or 30 μg nanoparticles in the original samples and run by electrophoresis at 160 V. For fluorescence scans, the gels were first imaged by an Amersham Typhoon NIR laser scanner (GE Healthcare Life Sciences). Subsequently, protein bands were stained by Coomassie Brilliant Blue (Imperial Protein Stain; ThermoFisher Scientific) and scanned on a Bio-Rad gel documentation system. Band intensity quantification was performed using the plot profile tool in Fiji/ImageJ.^45, 46^ Three independent experiments were performed for SDS-PAGE and representative results are shown.

### Protein identification on excised bands

The in-gel digestion of selected protein bands was performed as previously described.^47^ Prior to analysis by mass spectrometry, the tryptic peptides were micropurified using Empore SPE Disks of C18 octadecyl packed in 10 μl pipette tips.^48^ LC-MS/MS analyses were performed on an Eksigent nanoLC 415 system (SCIEX) connected to a TripleTOF 6600 mass spectrometer (SCIEX) equipped with a NanoSpray III source (AB SCIEX). The trypsin-digested samples were suspended in 0.1% formic acid, injected, trapped and desalted isocratically on a precolumn (ReproSil-Pur C18-AQ 3 μm resin, Dr. Maisch GmbH, Germany). The peptides were eluted and separated on a 15 cm analytical column (75 μm i.d.) packed with ReproSil-Pur C18-AQ 3 μm resin in a pulled emitter. Peptides were eluted at a flow rate of 250 nl/min using a 30 min gradient from 5% to 35% of solution B (0.1% formic acid, 100% acetonitrile). The collected MS files (.wiff) were converted to Mascot generic format (MGF) using the AB SCIEX MS Data Converter beta 1.1 (AB SCIEX). The generated peak lists were searched using an in-house Mascot search engine (Matrix Science). Search parameters were set to allow one missed trypsin cleavage site and propionamide as a fixed modification with peptide tolerance and MS/MS tolerance set to 20 ppm and 0.4 Da, respectively.

### Zebrafish

Zebrafish (*Danio rerio*) were bred and maintained following the Danish legislation under permit number 2017-15-0202-00098. Experiments were performed on 3 dpf embryos from a wild-type strain (AB) or established transgenic lines; *Tg(fli1a:EGFP)^y1^* and *Tg(kdrl:Hsa.HRAS-mCherry)^s916^* or *Tg(kdrl:mCherry-F)* for short were used as reporter lines for endothelial cells;^49–51^ *Tg(mpeg1:mCherry)^g123^* for embryonic macrophages;^52^ *Tg(tnfa:EGFP-F)^ump5^* for transcriptional activation of *tnfa.^52^*

### Intravital confocal laser scanning microscopy and image analysis

In the same manner as described previously,^24^ anesthetized zebrafish embryos (3 dpf) were embedded in 0.8% (w/v) low-melting-point agarose for IV microinjection with nominal 10 ng (3 × 10^7^ particles, 3 nl at 3.4 mg/ml) of nanoparticles along with sterile-filtered, endotoxin-free phenol red solution in DPBS (0.5% w/v, BioReagent; Sigma-Aldrich) as a loading dye/buffer. As a vehicle control, water (MilliQ, 18.2 MΩ) was used likewise diluted in the loading buffer. The injection dose of free FBS proteins was 2 ng, equivalent to the measured protein mass of FBS-PC nanoparticles. For M1-like polarization of macrophages, as previously performed,^24^ γ-irradiated LPS from *Escherichia coli* O111:B4 (BioXtra; Sigma-Aldrich) dissolved in PBS was IV-injected at the dose of 2.4 ng, which is 1,000,000-fold higher than the measured endotoxin level detected in FBS-PC nanoparticles. Injected embryos were observed non-invasively under a Zeiss LSM 780 upright confocal microscope (Carl Zeiss) with lasers at 405 nm (Pacific Blue), 488 nm (EGFP/FITC), 568 nm (mCherry, pHrodo Red), and 633 nm (Cy5). Objective lenses used were EC Plan-Neofluar 10x/0.3 M27 (Carl Zeiss) for tile scanning of the whole embryo and W Plan-Apochromat 40×/1.0 DIC M27 (Carl Zeiss) for live imaging. All images at the ROI were acquired at 2-μm intervals ensuring optical section overlap in each fluorescence channel to construct *z*-stack images and presented as the maximum intensity *z*-stack projections using Fiji/ImageJ.^45, 46^ Detailed descriptions of the 3D mask approach and image analysis method are fully presented in our previous report.^24^ Similarly to “sequestered nanoparticles” defined therein, “sequestered PC” was defined by thresholding of the PC signals at >20 times higher than the noise level typically picking only the immobilized clusters associated with cells. Likewise, “Total FI” is defined by mean fluorescence intensity multiplied by area (μm^2^) after thresholding to remove the background noise signals. For fluorescence ratio quantification, we used the 3D mask approach whereby the colocalization of Cy5 and pHrodo or Pacific Blue signals was determined at each focal plane of 2 μm-thick optical sections before maximum intensity projection of a stacked image. Briefly, as mathematical pre-processing for calculating a fluorescence ratio in each pixel by division, a value of 1 was assigned to all pixels in both fluorescence channels before applying a 3D mask specific for FBS PC (Figure 3, FBS-Cy5 as the mask for FBS-pHrodo) or nanoparticles (Figure 4, NP-Pacific Blue as the mask for FBS-Cy5). The masked, stacked images were then processed as maximum intensity *z*-projections and the fluorescence ratio of two channels was calculated in the image using a 32-bit floating-point mode. The resulting values were multiplied by a factor of 10,000 to convert the images back to the 16-bit range (0-65535), where the value of 10,000 equals to the fluorescence ratio of 1. To obtain a representative value for the fluorescence ratios, the median, rather than the mean, is reported. The 3D mask approach takes into account pixel-by-pixel colocalization of two fluorescent signals, but we could also obtain similar results when we simply calculated the fluorescence ratio using median values from each signal channel without considering colocalization. All intravital imaging experiments were repeated independently at least once more to confirm the reproducibility.

### Kinetic models and statistics

Non-linear curve fitting was performed in R (ver. 3.5.1) for results obtained from time-lapse imaging. All individual data points were fitted using an exponential decay model; *y* – *y_f_* + (*y*_0_ – *y_f_*)*e*^−*λt*^, where the measured value *y* starts at *y*_0_ and decreases towards *y_f_* as an exponential function of time *t* with the rate constant *λ*, or logistic/Gompertz models; 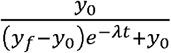 and 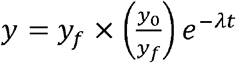, where the starting value *y*_0_ grows towards the plateau *y_f_* as an exponential function of time *t* with the rate constant *λ*. The fitted parameters are shown in Supporting Table 3. For parametric statistics, the input data were log-transformed to satisfy the assumption of normality. One-way analysis of variance (ANOVA) with Tukey’s HSD post-hoc comparisons were performed in R (ver. 3.5.1) following Levene’s test on the homogeneity of variances. Significant differences between the tested populations were determined as *α* = 0.05.

### Transmission electron microscopy

Ultrathin-sectioned specimens of nanoparticle-injected embryos (30 mpi) were prepared exactly the same manner as described previously in full detail,^24^ and examined under a Tecnai G2 Spirit TEM (FEI company, ThermoFisher Scientific) operating at 120 KeV.

## Supporting information

Movie 1 (Unmodified)

Movie 2 (FBS-PC)

Movie 3 (PC acidification)

Movie 4 (kdrl Water)

Movie 5 (kdrl Unmodified)

Movie 6 (kdrl PC)

Movie 7 (tnfa Water)

Movie 8 (tnfa Unmodified)

Movie 9 (tnfa Protein)

Movie 10 (tnfa PC)

Movie 11 (tnfa LPS)

## ASSOCIATED CONTENT

The authors declare no competing financial interest.

## Supporting Information

The Supporting Information is available free of charge on the ACS Publications website.

Supporting tables, figures and description of movie files (PDF)

Movie 1: Sequestration of unmodified nanoparticles at 3-30 mpi (AVI)

Movie 2: Sequestration of FBS-PC nanoparticles at 3-30 mpi (AVI)

Movie 3: Endolysosomal acidification of FBS-PC nanoparticles at 3-24 mpi (AVI)

Movie 4: Blood vessel integrity at 1-12 hpi, water control (AVI)

Movie 5: Blood vessel integrity at 1-12 hpi, unmodified nanoparticles (AVI)

Movie 6: Blood vessel integrity at 1-12 hpi, FBS-PC nanoparticles (AVI)

Movie 7: M1-like polarization of macrophages at 1-12 hpi, water control (AVI)

Movie 8: M1-like polarization of macrophages at 1-12 hpi, unmodified nanoparticles (AVI)

Movie 9: M1-like polarization of macrophages at 1-12 hpi, FBS protein control (AVI)

Movie 10: M1-like polarization of macrophages at 1-12 hpi, FBS-PC nanoparticles (AVI)

Movie 11: M1-like polarization of macrophages at 1-12 hpi, LPS control (AVI)

## AUTHOR INFORMATION

**Corresponding Author**

*.

## ACKNOWLEDGMENT

This work was financially supported by the grants R219-2016-327 and R324-2019-1644 from Lundbeck Foundation (Y.H.), the grant DFF-4181-00473 from Independent Research Fund Denmark | the Research Council for Natural Sciences (FNU) and the center grant CellPAT (DNRF135) from the Danish National Research Foundation Center (H.M-B. and D.S.S.). We gratefully acknowledge P. Engelmann from the University of Pécs, Hungary, for his advice on the manuscript. We also thank the fish facility at Aarhus University for zebrafish husbandry.

## For Table of Contents Only

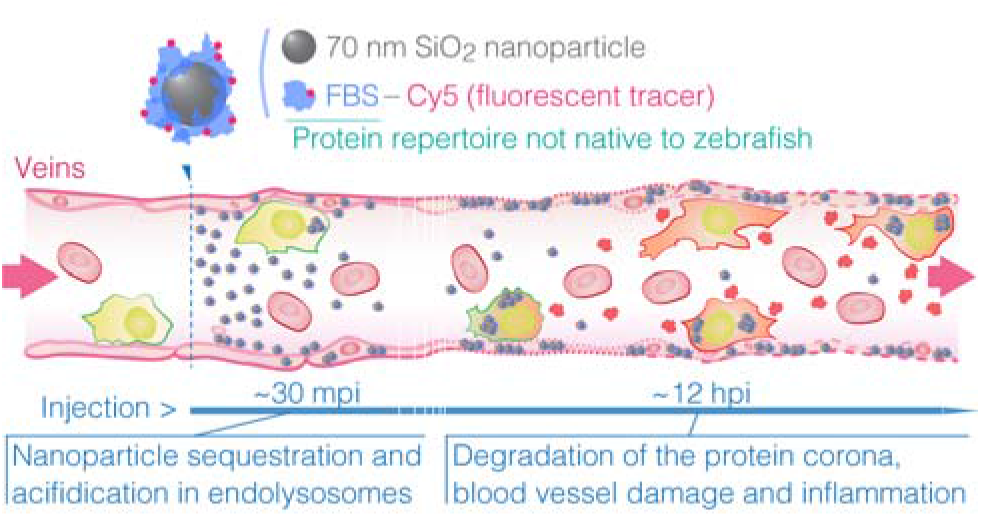

