## Supporting Information for "Tracing the *In Vivo* Fate of Nanoparticles with a “Non-Self” Biological Identity"

### Supporting Table 1

### Characterization summary.

| Nominal size | Surface | Fluorescent dye | Protein corona | Characterization data | |
|---|---|---|---|---|---|
| | | | | hydrodynamic size [a] (nm [PdI]) | ζ potential [b] (mV) |
| 70 nm | Plain | FITC | – | 74.0 ± 7.6 [0.042] | −20 ± 2 |
| | | | FBS | 103.4 ± 13.6 [0.069] | −19 ± 4 |
| 70 nm | Plain | Pacific Blue | – | 62.4 ± 13.3 [0.182] | −21 ± 3 |
| | | | FBS | 109.1 ± 23.2 [0.181] | −18 ± 2 |
| 70 nm | $NH_2$ | FITC | – | 74.3 ± 8.5 [0.052] | −15 ± 2 |
| | | | FBS | 126.1 ± 17.5 [0.077] | −17 ± 3 |
| 70 nm | COOH | FITC | – | 78.6 ± 5.7 [0.021] | −30 ± 6 |
| | | | FBS | 128.8 ± 23.6 [0.135] | −20 ± 3 |

[a] Dynamic light scattering; values are z-average ± PdI width obtained by the cumulant analysis.
[b] 10 mM sodium phosphate buffer (pH 7.4) as the dispersion medium. Values are mean ± SD of three measurements.

**Supporting Table 2**

**List of proteins identified in LC-MS/MS analysis of excised bands.**

| Band # (Mw) | Protein name (Gene name) | Symbol | Accession # | Mw (kDa) | Species | Protein score | Significant matches | Significant sequences[a] | Coverage % |
|---|---|---|---|---|---|---|---|---|---|
| 1 (72 kDa) | Ahsg protein (*alpha-2-HS-glycoprotein 1*) | Ahsg1 (Fetua) | Q5U3D8 | 50.1 | *Danio rerio* | 3127 | 113 | 5 (5) | 13 |
| | Zgc:153748 (*3'-phosphoadenosine 5'-phosphosulfate synthase 2a*) | Papss2a | A0JPF3 | 69.2 | *Danio rerio* | 2275 | 79 | 17 (15) | 34 |
| | Polyadenylate-binding protein 1A (*poly(A) binding protein, cytoplasmic 1a*) | Pabpc1a | F1QB54 | 71.0 | *Danio rerio* | 1521 | 51 | 19 (7) | 30 |
| 2 (15 kDa) | Hemoglobin subunit beta-2 (*hemoglobin, beta adult 2*) | Hbba2 | Q90485 | 16.4 | *Danio rerio* | 12571 | 359 | 10 (2) | 69 |
| | Hemoglobin, subunit beta-1 (*hemoglobin, beta adult 1*) | Hbba1 | Q90486 | 16.4 | *Danio rerio* | 12387 | 347 | 10 (2) | 69 |
| | Hemoglobin subunit alpha (*hemoglobin, alpha adult 1*) | Hbaa1 | Q90487 | 15.5 | *Danio rerio* | 4082 | 197 | 8 (4) | 52 |
| | Si:xx-by187g17.5 (*hemoglobin, alpha adult 2*) | Hbaa2 | Q5BJC7 | 15.4 | *Danio rerio* | 3634 | 130 | 4 (3) | 37 |
| | Apolipoprotein A-II (*apolipoprotein A-II*) | Apoa2 | B3DFP9 | 15.5 | *Danio rerio* | 3235 | 86 | 9 (9) | 43 |
| 3 (59 kDa) | ALB protein (*albumin*) | ALB | B0JYQ0 | 69.3 | *Bos taurus* | 9664 | 346 | 24 (2) | 29 |
| | Alpha-2-HS-glycoprotein (*alpha-2-HS-glycoprotein*) | AHSG (FETUA) | B0JYN6 | 38.4 | *Bos taurus* | 5009 | 123 | 9 (9) | 24 |
| | Alpha-1-antiproteinase (*alpha-1-antitrypsin*) | SERPINA1 | P34955 | 46.1 | *Bos taurus* | 4128 | 165 | 20 (20) | 38 |
| 4 (15 kDa) | Apolipoprotein A-II (*apolipoprotein A-II*) | APOA2 | P81644 | 11.2 | *Bos taurus* | 3446 | 166 | 4 (4) | 22 |

[a] Parentheses indicate the number of unique peptides.

**List of proteins identified in LC-MS/MS analysis of excised bands (continued).**

| Band # (Mw) | Protein name (Gene name) | Symbol | Accession # | Mw (kDa) | Species | Protein score | Significant matches | Significant sequences[a] | Coverage % |
|---|---|---|---|---|---|---|---|---|---|
| 5 (72 kDa) | Ahsg protein (*alpha-2-HS-glycoprotein 1*) | Ahsg1 (Fetua) | Q5U3D8 | 50.1 | *Danio rerio* | 3799 | 132 | 8 (8) | 19 |
| | Alpha-2-HS-glycoprotein (*alpha-2-HS-glycoprotein*) | AHSG (FETUA) | B0JYN6 | 38.4 | *Bos taurus* | 621[b] | 16 | 5 (5) | 18 |
| 6 (23 kDa) | Apolipoprotein A-I (*apolipoprotein A-I*) | APOA1 | P15497 | 30.3 | *Bos taurus* | 32078 | 1086 | 33 (33) | 83 |
| | Apolipoprotein A-Ib (*apolipoprotein A-Ib*) | Apoa1b | A0A0R4 IKF0 | 30.1 | *Danio rerio* | 4029 | 149 | 19 (19) | 57 |
| | Apolipoprotein A-I (*apolipoprotein A-I*) | Apoa1 | O42363 | 30.3 | *Danio rerio* | 2709 | 93 | 22 (22) | 62 |
| 7 (23 kDa) | Apolipoprotein A-I (*apolipoprotein A-I*) | APOA1 | P15497 | 30.3 | *Bos taurus* | 12244 | 496 | 28 (28) | 62 |
| | Apolipoprotein A-Ib (*apolipoprotein A-Ib*) | Apoa1b | A0A0R4 IKF0 | 30.1 | *Danio rerio* | 12222 | 468 | 16 (16) | 47 |
| | Apolipoprotein A-I (*apolipoprotein A-I*) | Apoa1 | O42363 | 30.3 | *Danio rerio* | 2221 | 80 | 23 (23) | 65 |

[a] Parentheses indicate the number of unique peptides.
[b] This protein may not represent the band excised but the score information is included for a comparison to the zebrafish orthologue.

**Supporting Table 3**

**Kinetic models used and fitted parameters (Exponential decay).**

| Model | Figure | RSE [a] (d.f.) | Fitted parameters | | |
|---|---|---|---|---|---|
| | | | $\lambda$ | $y_0$ | $y_f$ |
| Exponential decay $y = y_f + (y_0 - y_f)e^{-\lambda t}$ | Figure 2d (Unmodified) | 3710 (28) | 0.007 | 39010 | 0 |
| | Figure 2d (FBS-PC) | 667 (47) | 0.200 | 12360 | 1670 |
| | Figure 4f (Water) | 668 (222) | 0.020 | 9768 | 0 |
| | Figure 4f (Unmodified) | 705 (222) | 0.020 | 10260 | 0 |
| | Figure 4f (FBS-PC) | 846 (222) | 0.099 | 6957 | 0 |
| | Figure 4g (FBS-PC) | 403 (129) | 0.166 | 3990 | 0 |
| | SI Figure 10d (Unmodified) | 1060 (222) | 0.171 | 10820 | 1091 |
| | SI Figure 10f (FBS-PC) | 911 (129) | 0.173 | 8617 | 0 |
| | Figure 5b (FBS) | 238 (201) | 0.136 | 1317 | 687 |
| | Figure 5b (FBS-PC) | 414 (201) | 0.552 | 2206 | 1437 |
| | Figure 5b (LPS) | 333 (405) | 0.309 | 1888 | 1005 |
| | Figure 5c (inset) | 11470 (201) | 0.053 | 33100 | 0 |

[a] Residual standard error on (degrees of freedom); the square root of the residual sum of squares divided by the residual degrees of freedom.

**Kinetic models used and fitted parameters (Logistic/Gompertz growth).**

| Model | Figure | RSE [a] (d.f.) | Fitted parameters | | |
|---|---|---|---|---|---|
| | | | $\lambda$ | $y_0$ | $y_f$ |
| Gompertz growth $y = y_f \times \left(\frac{y_0}{y_f}\right)^{e^{-\lambda t}}$ | Figure 2e (Unmodified) | 508 (27) | 0.037 | 1273 | 4000 [b] |
| | Figure 2f (FBS-PC) | 574 (47) | 0.053 | 303 | 10970 |
| | Figure 3c (Ratio) | 0.18 (21) | 0.299 | 0.46 | 2.58 |
| | SI Figure 10e (Unmodified) | 806 (222) | 0.826 | 290 | 1855 |
| Logistic growth $y = y_f \times \frac{y_0}{(y_f - y_0)e^{-\lambda t} + y_0}$ | Figure 5c (All MΦ) | 3.48 (201) | 0.162 | 1.70 | 100 [b] |
| | Figure 5c (M1-like MΦ) | 1.30 (201) | 0.207 | 0.48 | 20.6 |
| | Figure 5d (FBS-PC) | 15.83 (201) | 0.752 | 5.33 | 55.7 |
| | Figure 5d (LPS) | 10.15 (405) | 1.138 | 0.90 | 46.6 |

[a] Residual standard error on (degrees of freedom); the square root of the residual sum of squares divided by the residual degrees of freedom.
[b] The fitted value corresponds to the upper constraint manually defined in the port algorithm.

**Supporting Figure 1**

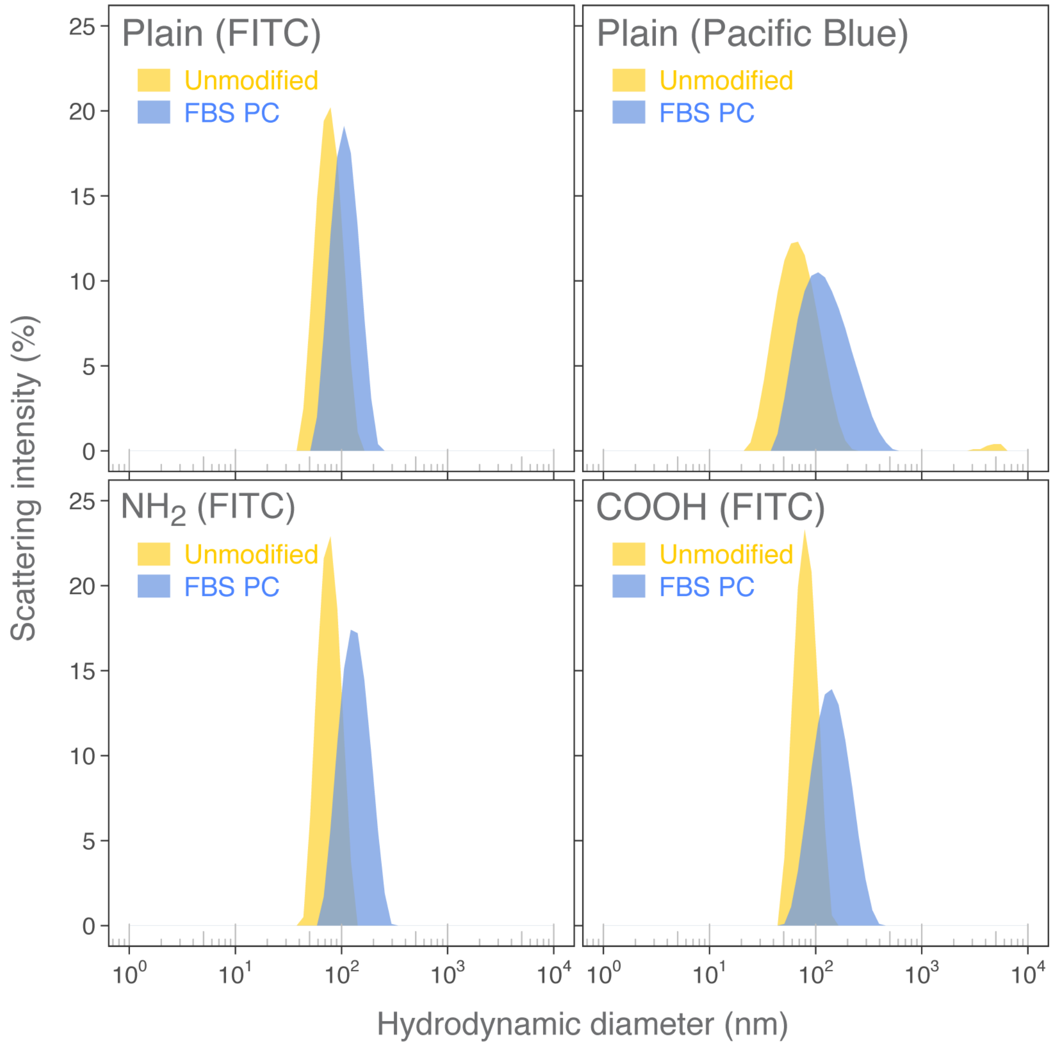

**Supporting Figure 1.** DLS analysis on the hydrodynamic size distributions. Size distributions of 70 nm $SiO_2$ nanoparticles with or without pre-formed FBS PC, fitted by the CONTIN algorithm. Three different surfaces (Plain, $NH_2$ and COOH) and two different dyes (FITC and Pacific Blue) used in this study were analyzed.

**Supporting Figure 2**

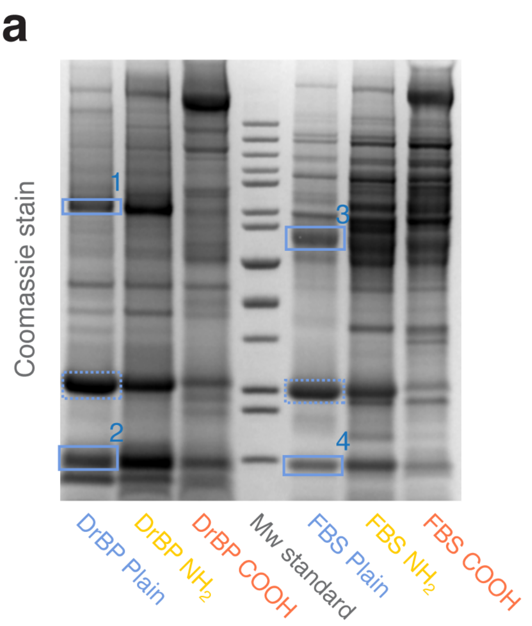

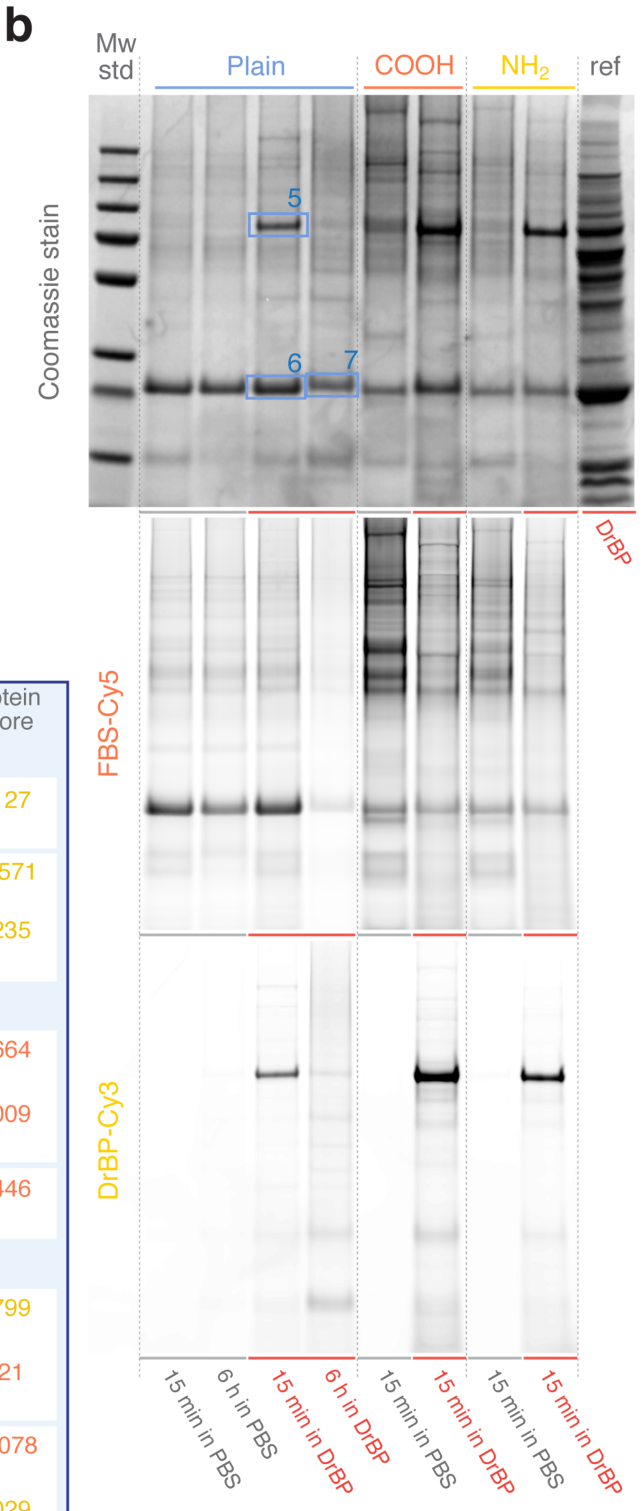

**c**

| band | protein | species origin | protein score |
|---|---|---|---|
| DrBP as the protein source | | | |
| 1 | alpha-2-HS-glycoprotein 1 (Ahsg1; aka Fetua) | *Danio rerio* | 3127 |
| 2 | hemoglobin subunit (Hb) | *Danio rerio* | 12571 |
| | apolipoprotein A-II (Apoa2) | *Danio rerio* | 3235 |
| FBS as the protein source | | | |
| 3 | albumin (ALB) | *Bos taurus* | 9664 |
| | alpha-2-HS-glycoprotein (AHSG; aka FETUA) | *Bos taurus* | 5009 |
| 4 | apolipoprotein A-II (APOA2) | *Bos taurus* | 3446 |
| FBS PC + 15 min in DrBP | | | |
| 5 | alpha-2-HS-glycoprotein 1 (Ahsg1; aka Fetua) | *Danio rerio* | 3799 |
| | alpha-2-HS-glycoprotein (AHSG; aka FETUA) | *Bos taurus* | 621 |
| 6 | apolipoprotein A-I (APOA1) | *Bos taurus* | 32078 |
| | apolipoprotein A-Ib (Apoa1b) | *Danio rerio* | 4029 |
| FBS PC + 6 h in DrBP | | | |
| 7 | apolipoprotein A-I (APOA1) | *Bos taurus* | 12244 |
| | apolipoprotein A-Ib (Apoa1b) | *Danio rerio* | 12222 |

**Supporting Figure 2.** Original gel images and protein identification. (a) Original gel images and (b) fluorescence scans used for Figure 1c-e additionally showing the results from two different surface functionalizations (-$NH_2$ and -COOH) and at the 6 h time point for the Plain surface. Note that the nominal mass of nanoparticles from which the PC was stripped is different between the gels in a (150 $\mu$g) and b (30 $\mu$g). (c) MS/MS protein identification of the selected bands designated in (a) and (b). The bands designated with dashed line rectangles in (a) were excluded from analysis since we have previously identified them as Apoa1b/APOA1 in similar experiments[1] and in this study after 15 min/6 h incubation in DrBP. Protein entries in yellow and orange are from protein databases for *Danio rerio* and *Bos taurus*, respectively. See Supporting Table 2 for details of the identified proteins.

**Supporting Figure 3**

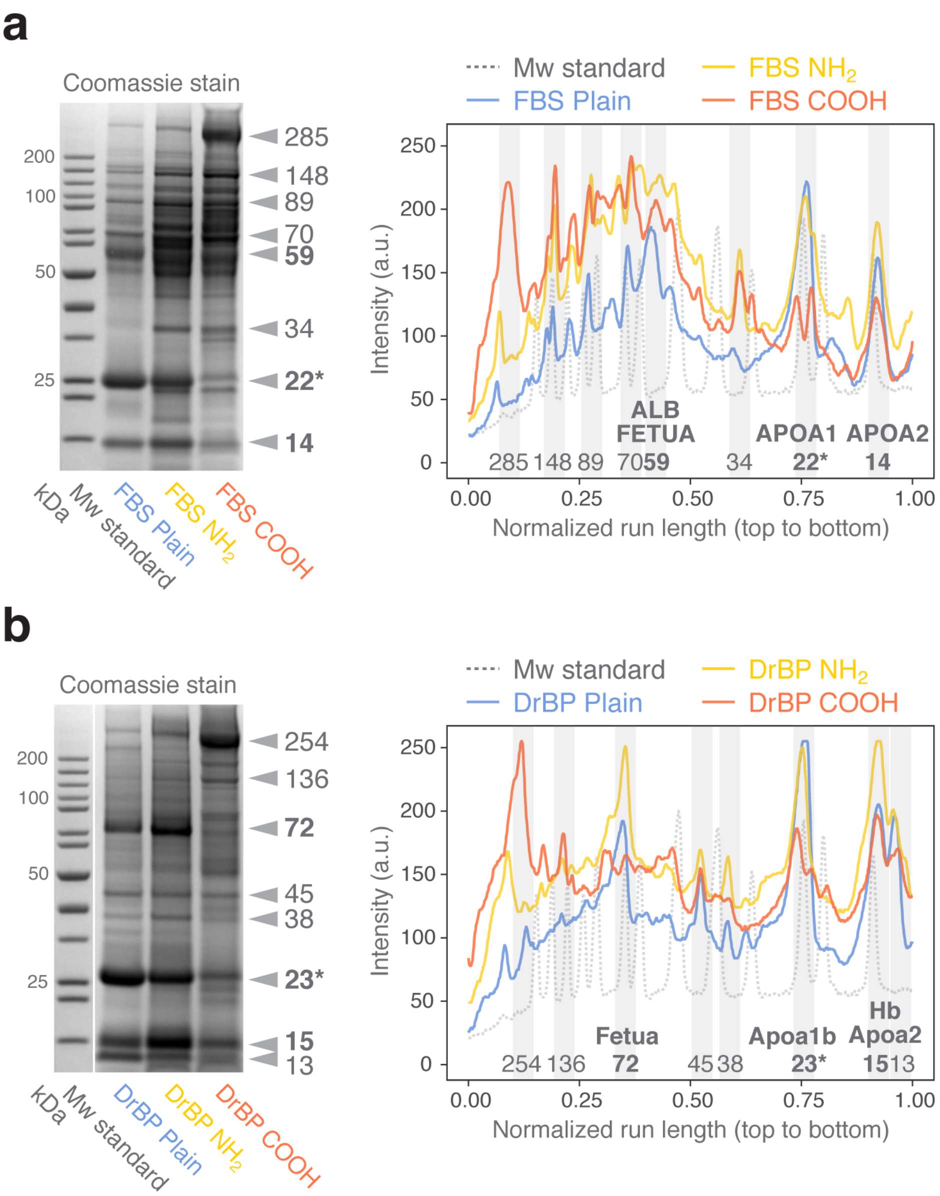

**Supporting Figure 3.** Comparison of SDS-PAGE profiles for different surface functionalizations. (a,b) SDS-PAGE profiles for (a) FBS PC and (b) DrBP PC. Major protein bands common to all surface types are indicated as arrowheads (gel images) and as gray-shaded areas (profile plots) along with molecular weights estimated from the molecular weight standards and, where available, the protein names identified (see Supporting Table 2). Asterisks denote the proteins identified in our previous report.[1] a.u., arbitrary unit.

**Supporting Figure 4**

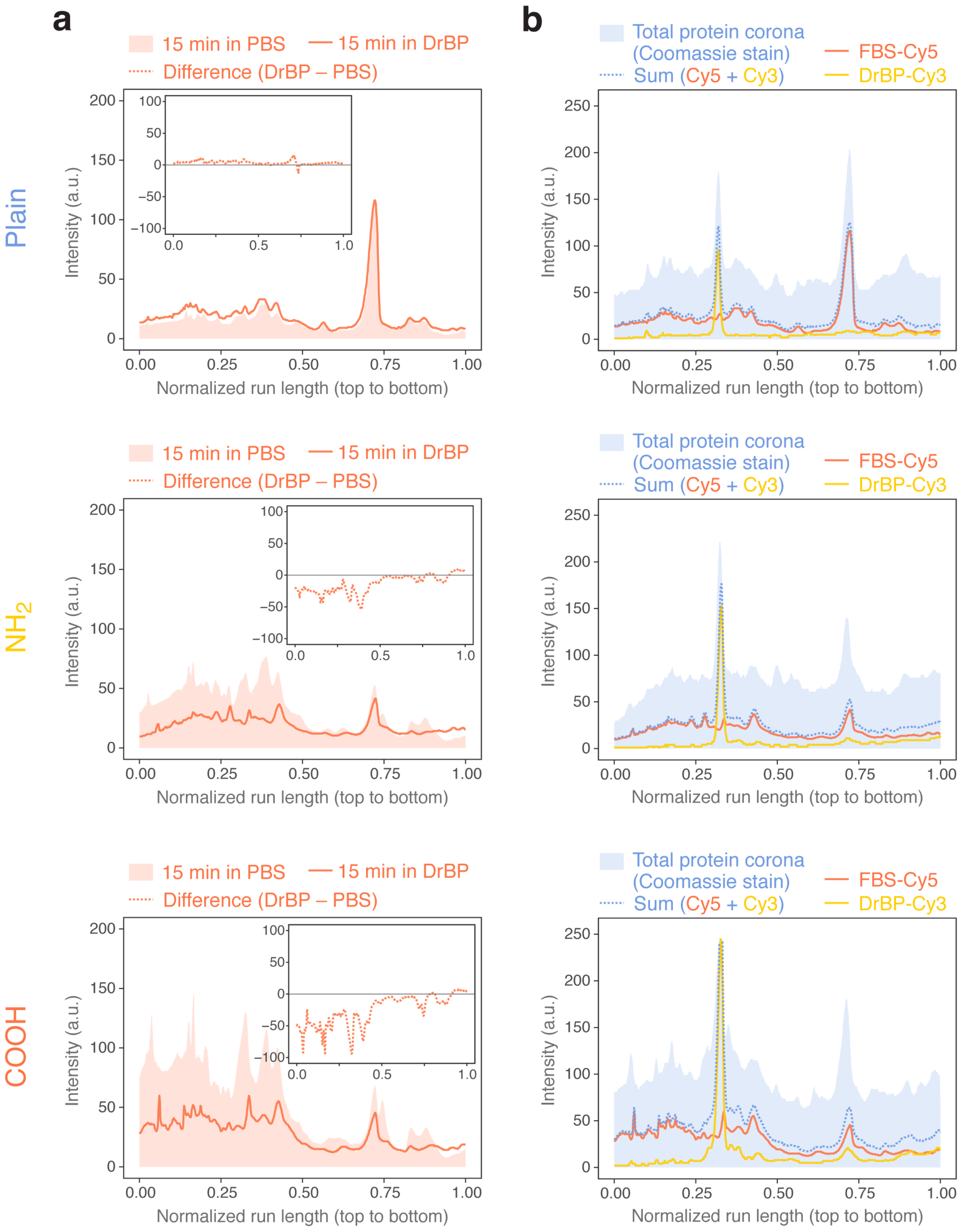

**Supporting Figure 4.** PC retention profiles for different surface functionalizations. (a) PC retention profiles for FBS-Cy5 following 15 min incubation in PBS or DrBP. The difference is plotted in the inset, showing a major decrease of higher molecular weight proteins for $NH_2$ and COOH. (b) The modified PC compositions after 15 min incubation in DrBP. Dashed line is the sum of FBS-Cy5 and DrBP-Cy3 signals that generally follows the peak positions identified by Coomassie Brilliant Blue staining of the total PC. See Supporting Figure 2b for the fluorescence scans. a.u., arbitrary unit.

**Supporting Figure 5**

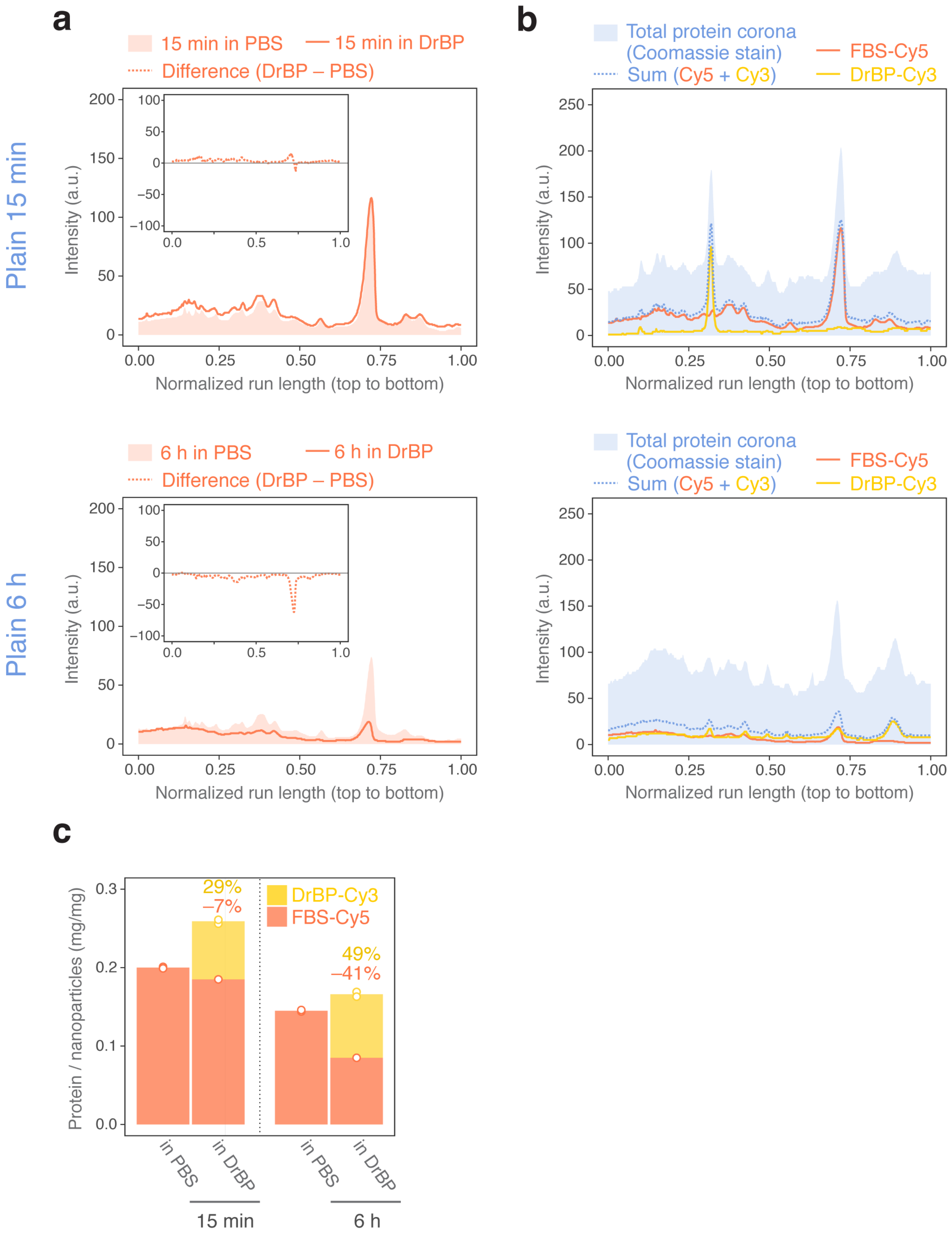

**Supporting Figure 5.** PC retention profiles at two time points. (a) PC retention profiles for FBS-Cy5 following 15 min and 6 h incubation in PBS or DrBP. The difference is plotted in the inset, showing a major decrease of the band representing APOA1 at 6 h. (b) The modified PC compositions after 15 min and 6 h incubation in DrBP. Dashed line is the sum of FBS-Cy5 and DrBP-Cy3 signals that generally follows the peak positions identified by Coomassie Brilliant Blue staining of the total PC. Note the decrease of the band representing Fetua (DrBP-Cy3) and nearly equal contribution of APOA1 (FBS-Cy5) and Apoa1b (DrBP-Cy3) for the band corresponding to Apoa1b indicating competitive replacement of FBS by DrBP at 6 h. See Supporting Figure 2b,c for the fluorescence scans and protein scores for APOA1 *versus* Apoa1b. a.u., arbitrary unit. (c) Fluorimetry-based quantification of FBS-Cy5 and DrBP-Cy3 in the PC. The values shown above the columns are the percentage decrease of FBS as compared to "in PBS" and the fraction of DrBP in the total PC. Columns represent the mean of two measurements (empty points).

### Supporting Figure 6

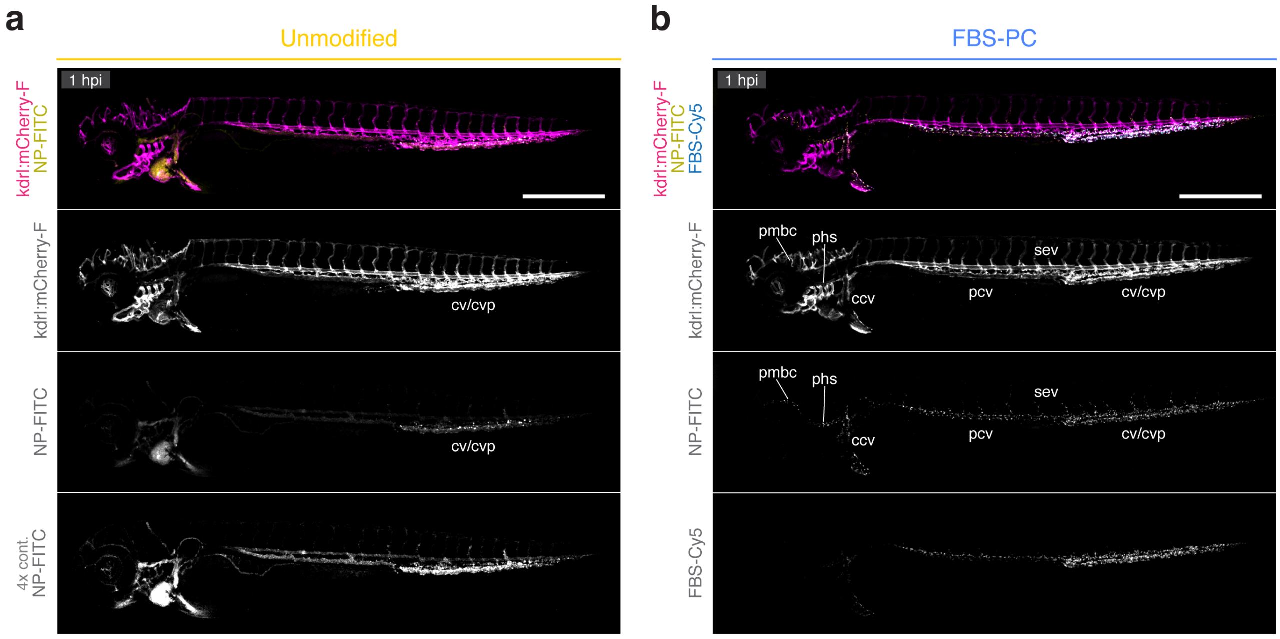

**Supporting Figure 6.** Biodistribution mapping of IV-injected nanoparticles. (a,b) *Tg(kdrl:mCherry-F)* embryos at 3 dpf were injected with FITC-labeled $SiO_2$ nanoparticles (NP-FITC) with or without pre-formed FBS PC (FBS-Cy5) and imaged at 1 hpi. Representative tiled images depict the blood vessel networks (*kdrl:mCherry-F*) and biodistribution of (a) unmodified and (b) FBS-PC nanoparticles. The NP-FITC signals of the bottom panel in a are multiplied by 4-fold to aid visualization of blood-borne nanoparticles that remain circulating in the bloodstream. Anatomical annotations are indicated where the nanoparticles are sequestered. Anterior left, dorsal top. Scale bars, 500 $\mu$m. pmbc, primordial midbrain channel. phs, primary head sinus. ccv, common cardinal vein. pcv, posterior caudal vein. cv, caudal vein. cvp, caudal vein plexus. sev, intersegmental veins.

**Supporting Figure 7**

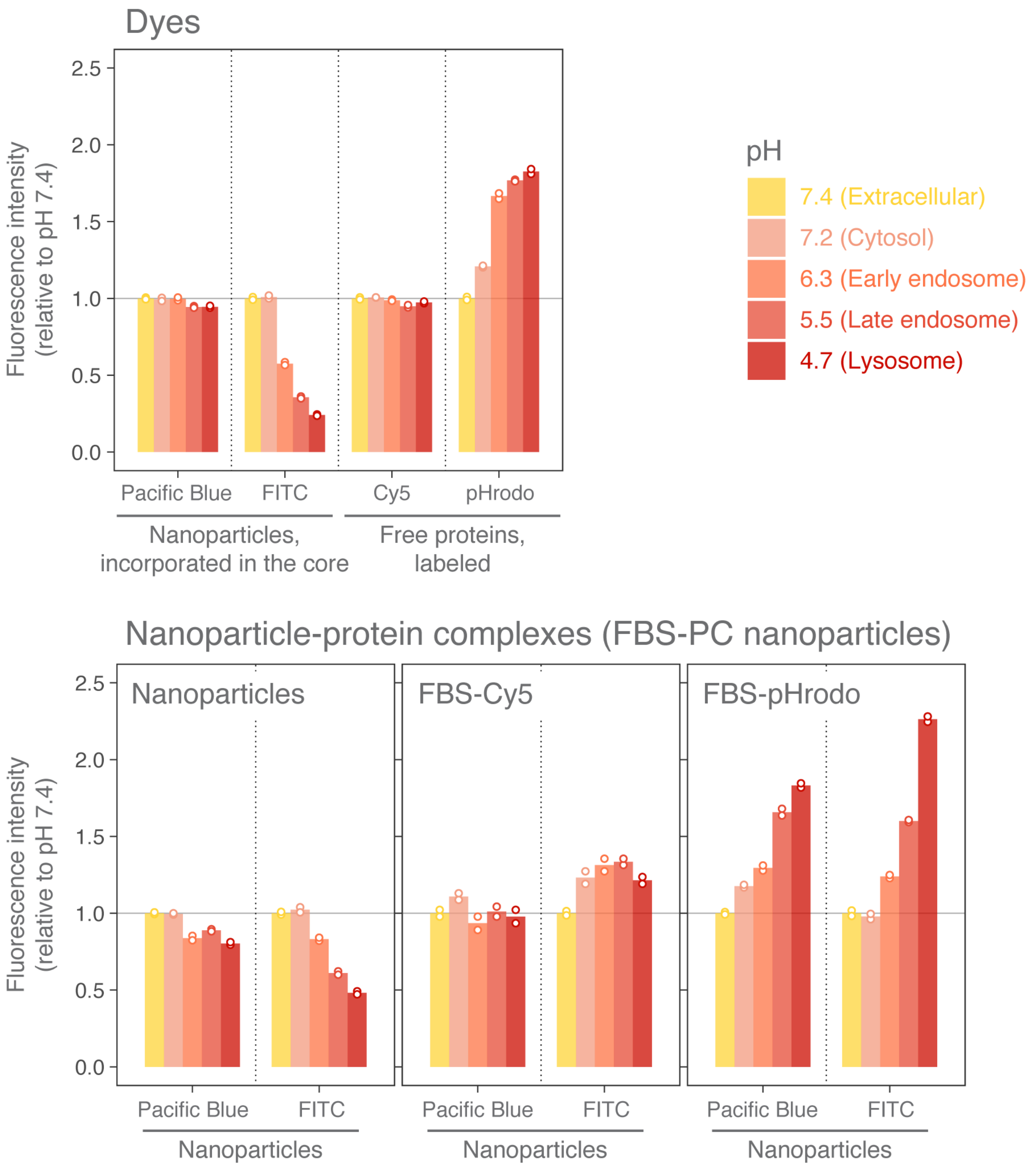

**Supporting Figure 7.** Effects of pH on the fluorescence intensity of FBS-PC nanoparticles. Fluorescence intensities of Pacific Blue and FITC nanoparticles, free proteins labeled with Cy5 or pHrodo, and the nanoparticle-protein complexes (FBS-PC nanoparticles) labeled with the mixture of Cy5 and pHrodo were measured by fluorimetry at varying pH relevant for intracellular environments. Columns represent the mean of two measurements (empty points). Phosphate-citrate buffer was used to achieve pH of choice while retaining the buffering capacity. DLS was performed on the nanoparticles at pH ranges between 4 to 7.4 and confirmed no pH-induced agglomeration.

### Supporting Figure 8

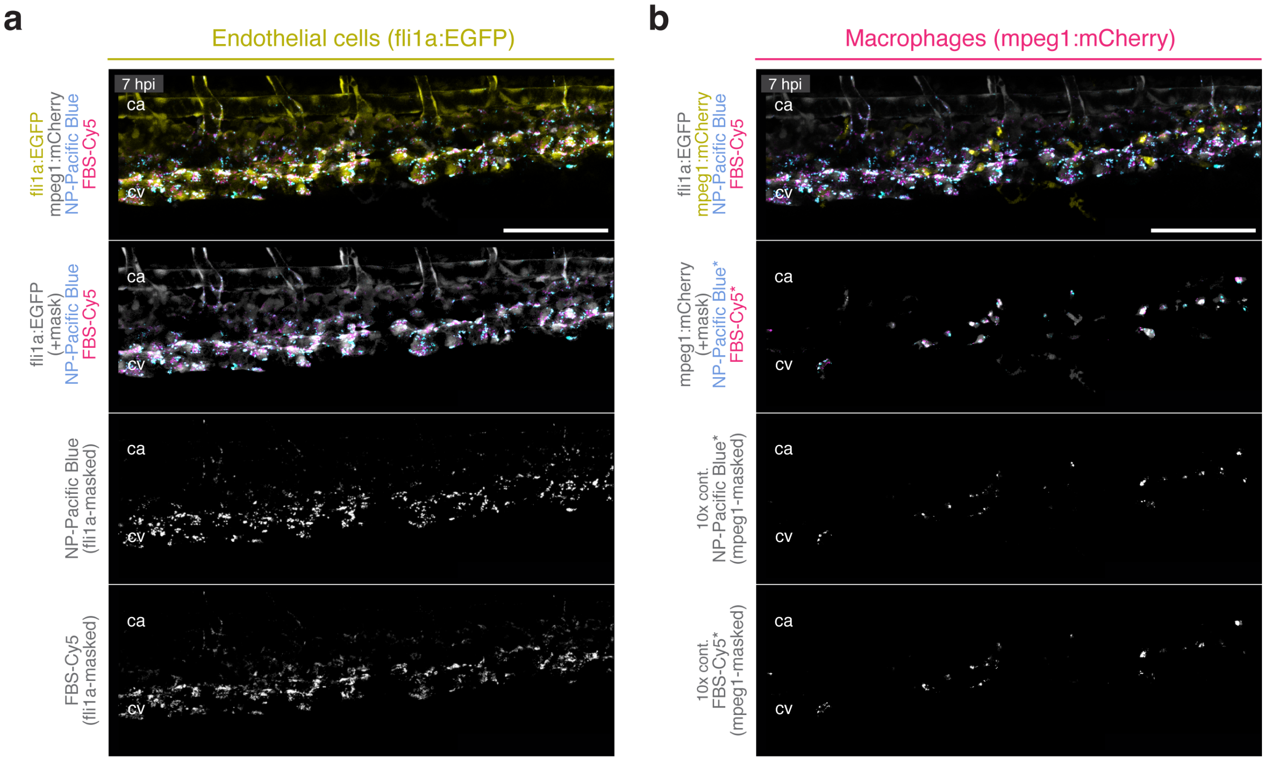

**Supporting Figure 8.** The 3D mask approach to analyze cell type-specific signals. (a,b) *Tg(fli1a:EGFP); Tg(mpeg1:mCherry)* embryos at 3 dpf were injected with Pacific Blue-labeled $SiO_2$ nanoparticles (NP-Pacific Blue) with pre-formed FBS PC (FBS-Cy5) and imaged at 7 hpi. Representative tiled images showing (a) EC- and (c) macrophage-specific signals for NP-Pacific Blue and FBS-Cy5. Asterisks in (b) indicate enhanced contrast (10-fold) applied to nanoparticle and FBS-Cy5 signals to aid visualization. Anterior left, dorsal top. Scale bars, 250 $\mu$m.

### Supporting Figure 9

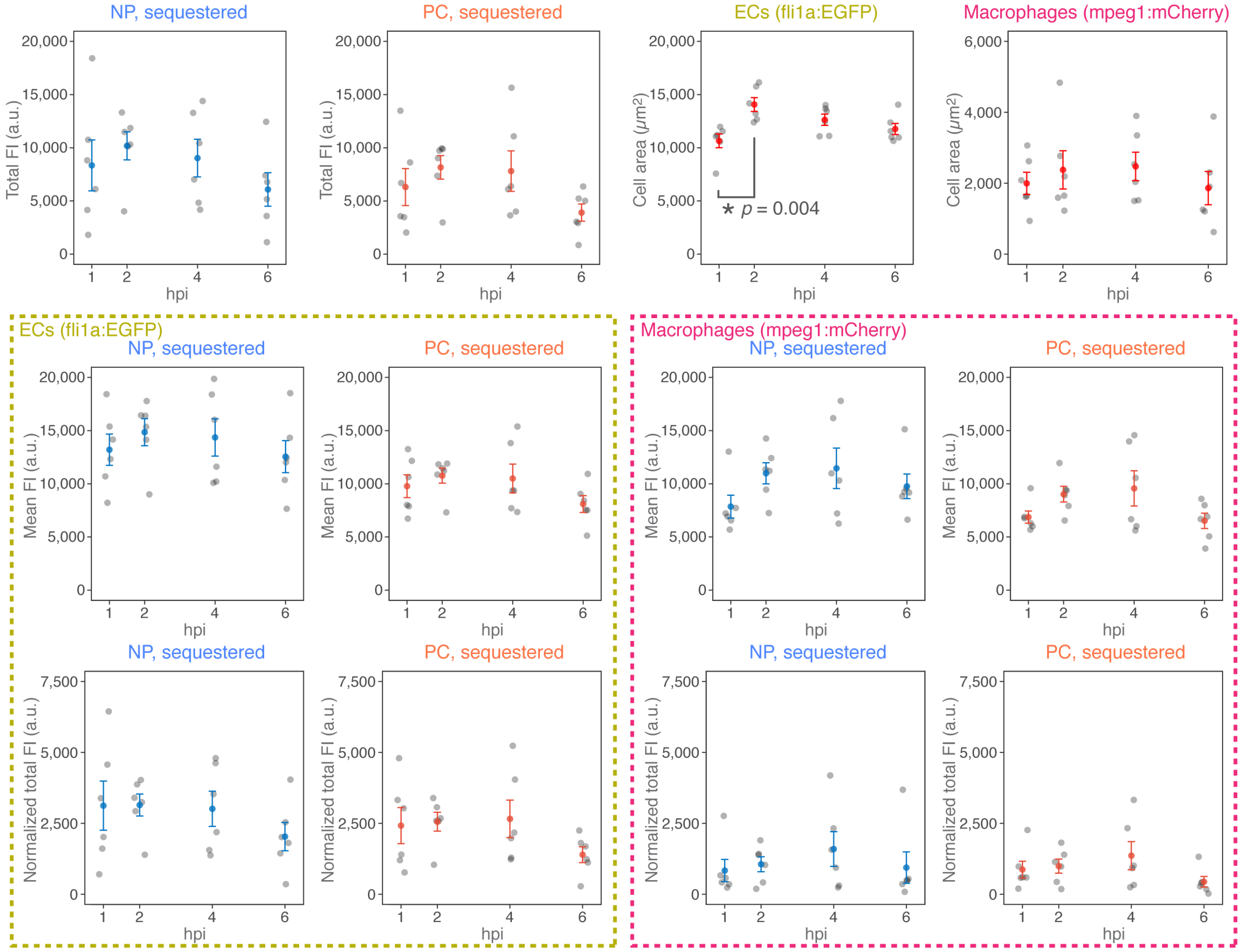

**Supporting Figure 9.** Supplementary image analysis results for Figure 4a-d. *Tg(fli1a:EGFP); Tg(mpeg1:mCherry)* embryos at 3 dpf were injected with Pacific Blue-labeled $SiO_2$ nanoparticles with pre-formed FBS PC for each time point of 1, 2, 4 and 6 hpi and imaged independently. Values are plotted as individual embryos (gray points, $n = 6$) and the mean ± SE (blue/orange/red-colored). Significant differences were tested by one-way ANOVA with Tukey's HSD post-hoc comparisons (degrees of freedom = 20). "Normalized total FI" is total FI divided by the cell area (ECs or macrophages). a.u., arbitrary unit. FI, fluorescence intensity.

**Supporting Figure 10**

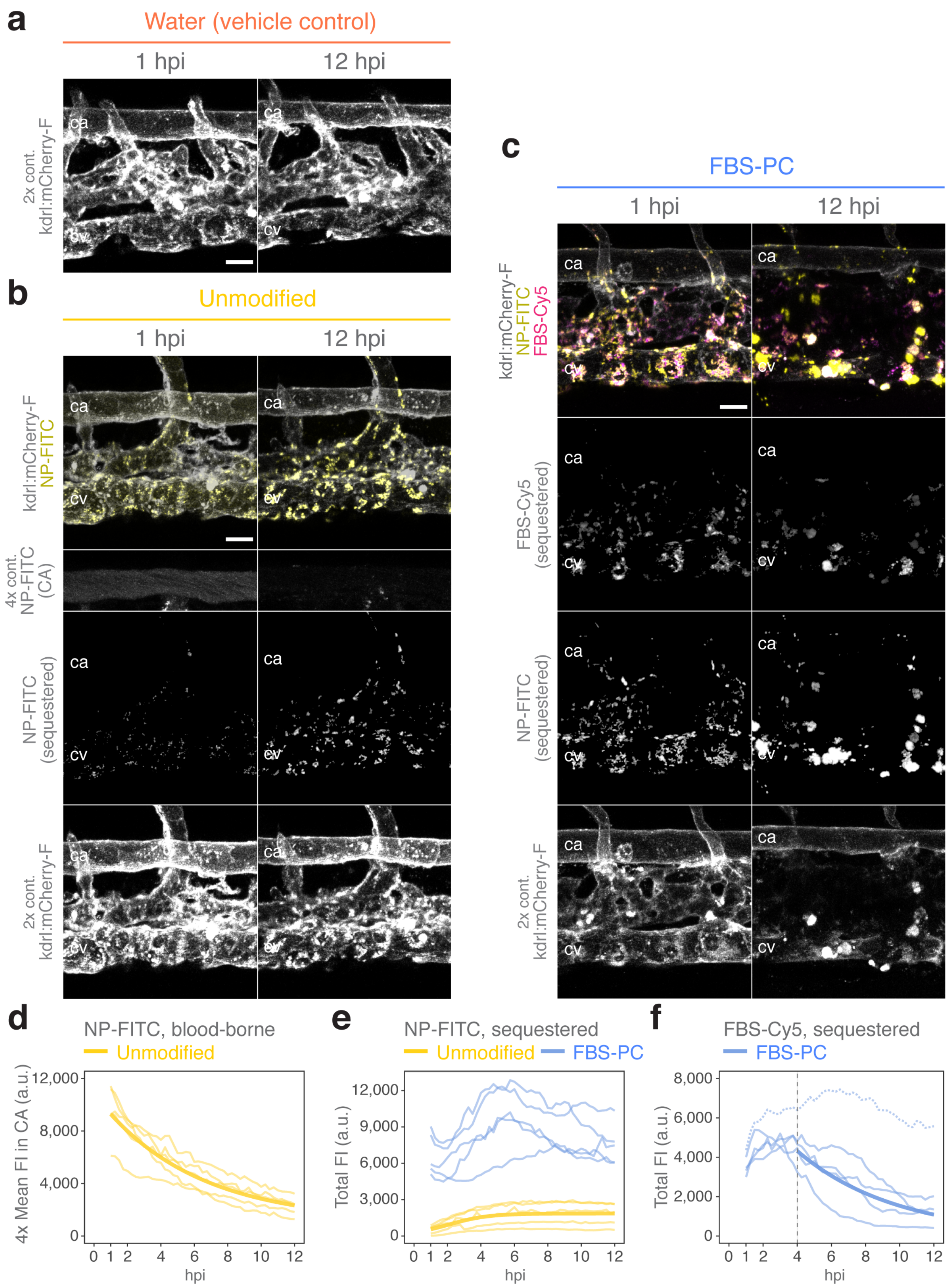

**Supporting Figure 10.** Supplementary image analysis results for Figure 4e-g. (a-f) *Tg(kdrl:mCherry-F)* embryos at 3 dpf were injected with FITC-labeled $SiO_2$ nanoparticles (NP-FITC) with or without pre-formed FBS PC (FBS-Cy5) and imaged every 15 min starting at 1 hpi. Representative images showing (a,b) the intact blood vessels, (b) blood clearance of unmodified nanoparticles, and (c) loss of fluorescence signals from scavenger ECs and sequestered FBS-PC nanoparticles both as NP-FITC and FBS-Cy5. The mCherry and NP-FITC (in CA) signals are multiplied by 2-fold and 4-fold, respectively, to aid visualization. Anterior left, dorsal top. Scale bars, 20 $\mu$m. Kinetics for (d) clearance of blood-borne nanoparticles and sequestration of (e) nanoparticles and (f) FBS PC are plotted as individual embryos (thin lines, $n = 5$) and, where possible, curve-fitted (thick lines). Note that the NP-FITC fluorescence is quenched after endolysosomal sequestration, resulting in the signal reduction over time even though the blood vessels are intact. The curve fitting in (f) was performed after excluding an outlier (the dashed line) and between 4-12 hpi where steady decreases are observed. a.u., arbitrary unit. FI, fluorescence intensity. See Movies 4-6 for the time-lapse sequences.

**Descriptions of movie files**

**Movie 1.** Sequestration of unmodified nanoparticles at 3-30 mpi. *Tg(kdrl:mCherry-F)* embryos at 3 dpf were injected with FITC-labeled $SiO_2$ nanoparticles (NP-FITC) without pre-formed FBS PC and imaged every 3 min. Representative movies showing sequestration of unmodified nanoparticles. The huge clusters of nanoparticles that move around are likely those associated with macrophages.[2] Left panel, merged channel for total signals: *kdrl:mCherry* in gray and NP-FITC in yellow. Right panel, single channel for sequestered nanoparticles: NP-FITC in gray. Anterior left, dorsal top.

**Movie 2.** Sequestration of FBS-PC nanoparticles at 3-30 mpi. *Tg(kdrl:mCherry-F)* embryos at 3 dpf were injected with FITC-labeled $SiO_2$ nanoparticles (NP-FITC) with pre-formed FBS PC (FBS-Cy5) and imaged every 3 min. Representative movies showing sequestration of FBS-PC nanoparticles. Left panels, merged channels for total signals: *kdrl:mCherry* in gray, NP-FITC in yellow and FBS-Cy5 in magenta, (top) included or (bottom) excluded. Right panels, single channels for sequestered nanoparticles: (bottom) NP-FITC in gray and (top) FBS-Cy5 in gray. Anterior left, dorsal top.

**Movie 3.** Endolysosomal acidification of FBS-PC nanoparticles at 3-24 mpi. Wild-type embryos at 3 dpf were injected with FITC-labeled $SiO_2$ nanoparticles with pre-formed FBS PC (FBS-Cy5 and FBS-pHrodo) and imaged every 3 min. Representative movies showing sequestration of dual-labeled FBS PC and the fluorescence ratios, 0-2 scaled. Left panel, merged channel for total signals: FBS-Cy5 in magenta and FBS-pHrodo in cyan. Right panel, fluorescence ratios for sequestered FBS PC: 0-2 scaled from purple/dark to yellow/white. Anterior left, dorsal top.

**Movie 4.** Blood vessel integrity at 1-12 hpi, water control. *Tg(kdrl:mCherry-F)* embryos at 3 dpf were injected with a vehicle control (water) and imaged every 15 min starting at 1 hpi. Representative movies showing intact blood vessels over time. The mCherry signals are multiplied by 2-fold to aid visualization. Single channel images for total signals: *kdrl:mCherry* in gray (right panel) with and (left panel) without the 2-fold enhanced contrast. Anterior left, dorsal top.

**Movie 5.** Blood vessel integrity at 1-12 hpi, unmodified nanoparticles. *Tg(kdrl:mCherry-F)* embryos at 3 dpf were injected FITC-labeled $SiO_2$ nanoparticles (NP-FITC) without pre-formed FBS PC and imaged every 15 min starting at 1 hpi. Representative movies showing sequestration of unmodified nanoparticles. Left/top panel, merged channel for total signals: *kdrl:mCherry* in gray and NP-FITC in yellow. Left/bottom panel, single channel for sequestered nanoparticles: NP-FITC in gray. Right/top panel, single channel for total signals: *kdrl:mCherry* in gray (2-fold enhanced contrast). Right/bottom panel, single channel for all nanoparticles: NP-FITC in gray (4-fold enhanced contrast to aid visualization of blood-borne nanoparticles). Anterior left, dorsal top.

**Movie 6.** Blood vessel integrity at 1-12 hpi, FBS-PC nanoparticles. *Tg(kdrl:mCherry-F)* embryos at 3 dpf were injected with FITC-labeled $SiO_2$ nanoparticles (NP-FITC) with pre-formed FBS PC (FBS-Cy5) and imaged every 15 min starting at 1 hpi. Representative images showing loss of fluorescence signals from scavenger ECs and sequestered FBS-PC nanoparticles. Left/top panel, merged channel for total signals: *kdrl:mCherry* in gray, NP-FITC in yellow and FBS-Cy5 in magenta. Left/bottom panel, single channel for sequestered nanoparticles: NP-FITC in gray. Right/top panel, single channel for total signals: *kdrl:mCherry* in gray (2-fold enhanced contrast). Right/bottom panel, single channel for sequestered FBS PC: FBS-Cy5 in gray. Anterior left, dorsal top.

**Movie 7.** M1-like polarization of macrophages at 1-12 hpi, water control. *Tg(mpeg1:mCherry); Tg(tnfa:GFP-F)* embryos at 3 dpf were injected with a vehicle control (water) and imaged every 20 min starting from 1 hpi. Representative movies showing no *tnfa* induction over time. Left panel, merged channel for total signals: *mpeg1:mCherry* in gray and *tnfa:EGFP-F* in yellow. Right panel, merged channel for macrophage-masked signals: *mpeg1:mCherry* in gray and *tnfa:EGFP-F* in yellow. Anterior left, dorsal top.

**Movie 8.** M1-like polarization of macrophages at 1-12 hpi, unmodified nanoparticles. *Tg(mpeg1:mCherry); Tg(tnfa:EGFP-F)* embryos at 3 dpf were injected with Pacific Blue-labeled $SiO_2$ nanoparticles (NP-Pacific Blue) without pre-formed FBS PC and imaged every 20 min starting from 1 hpi. Representative movies showing sequestration of unmodified nanoparticles and no *tnfa* induction over time. Note that NP-Pacific Blue is prone to photobleaching and the overall fluorescence intensity decreases over time. Left/top panel, merged channel for total signals: *mpeg1:mCherry* in gray, *tnfa:EGFP-F* in yellow and NP-Pacific Blue in cyan. Right/top panel, merged channel for macrophage-masked signals: *mpeg1:mCherry* in gray, *tnfa:EGFP-F* in yellow and NP-Pacific Blue in cyan. Left/bottom panel, merged channel for macrophage-masked signals: *mpeg1:mCherry* in gray and *tnfa:EGFP-F* in yellow. Right/bottom panel, merged channel for macrophage-masked signals: *mpeg1:mCherry* in gray and NP-Pacific Blue in cyan. Anterior left, dorsal top.

**Movie 9.** M1-like polarization of macrophages at 1-12 hpi, FBS protein control. *Tg(mpeg1:mCherry); Tg(tnfa:EGFP-F)* embryos at 3 dpf were injected with FBS proteins only (at the equivalent protein mass of FBS-PC nanoparticles) and imaged every 20 min starting from 1 hpi. Representative movies showing no sequestration of FBS proteins (FBS-Cy5) and no *tnfa* induction over time. Left/top panel, merged channel for total signals: *mpeg1:mCherry* in gray, *tnfa:EGFP-F* in yellow and FBS-Cy5 in magenta. Right/top panel, merged channel for macrophage-masked signals: *mpeg1:mCherry* in gray, *tnfa:EGFP-F* in yellow and FBS-Cy5 in magenta. Left/bottom panel, merged channel for macrophage-masked signals: *mpeg1:mCherry* in gray and *tnfa:EGFP-F* in yellow. Right/bottom panel, merged channel for macrophage-masked signals: *mpeg1:mCherry* in gray and FBS-Cy5 in magenta. Anterior left, dorsal top.

**Movie 10.** M1-like polarization of macrophages at 1-12 hpi, FBS-PC nanoparticles. *Tg(mpeg1:mCherry); Tg(tnfa:EGFP-F)* embryos at 3 dpf were injected with Pacific Blue-labeled $SiO_2$ nanoparticles (NP-Pacific Blue) with pre-formed FBS PC (FBS-Cy5) and imaged every 20 min starting from 1 hpi. Representative movies showing sequestration of FBS-PC nanoparticles and *tnfa* induction over time. Note that NP-Pacific Blue is prone to photobleaching and thus FBS-Cy5 is shown instead as a proxy for both components of FBS-PC nanoparticles. Left/top panel, merged channel for total signals: *mpeg1:mCherry* in gray, *tnfa:EGFP-F* in yellow and FBS-Cy5 in cyan. Right/top panel, merged channel for macrophage-masked signals: *mpeg1:mCherry* in gray, *tnfa:EGFP-F* in yellow and FBS-Cy5 in cyan. Left/bottom panel, merged channel for macrophage-masked signals: *mpeg1:mCherry* in gray and *tnfa:EGFP-F* in yellow. Right/bottom panel, merged channel for macrophage-masked signals: *mpeg1:mCherry* in gray and FBS-Cy5 in cyan. Anterior left, dorsal top.

**Movie 11.** M1-like polarization of macrophages at 1-12 hpi, LPS control. *Tg(mpeg1:mCherry); Tg(tnfa:EGFP-F)* embryos at 3 dpf were injected with a positive control (LPS) and imaged every 20 min starting from 1 hpi. Representative movies showing *tnfa* induction over time. Left panel, merged channel for total signals: *mpeg1:mCherry* in gray and *tnfa:EGFP-F* in yellow. Right panel, merged channel for macrophage-masked signals: *mpeg1:mCherry* in gray and *tnfa:EGFP-F* in yellow. Anterior left, dorsal top.
